# Forest edge landscape context affects mosquito community composition and risk of pathogen emergence

**DOI:** 10.1101/2024.04.30.591911

**Authors:** Adam Hendy, Nelson Ferreira Fé, Igor Pedrosa, André Girão, Taly Nayandra Figueira dos Santos, Claudia Reis Mendonça, José Tenaçol Andes Júnior, Flamarion Prado Assunção, Edson Rodrigues Costa, Vincent Sluydts, Marcelo Gordo, Vera Margarete Scarpassa, Michaela Buenemann, Marcus Vinícius Guimarães de Lacerda, Maria Paula Gomes Mourão, Nikos Vasilakis, Kathryn A. Hanley

## Abstract

Forest edges, where humans, mosquitoes, and wildlife interact, may serve as a nexus for zoonotic arbovirus exchange. Although often treated as uniform interfaces, the landscape context of edge habitats can greatly impact ecological interactions. Here, we investigated how the landscape context of forest edges shapes mosquito community structure in an Amazon rainforest reserve near the city of Manaus, Brazil, using hand-nets to sample mosquitoes at three distinct forest edge types. Sampling sites were situated at edges bordering urban land cover, rural land cover, and natural treefall gaps, while sites in continuous forest served as controls. Community composition differed substantially among edge types, with rural edges supporting the highest species diversity. Rural edges also provided suitable habitat for forest specialists, including key sylvatic vectors, of which *Haemagogus janthinomys* was the most abundant species sampled overall. Our findings emphasize the importance of landscape context in assessing pathogen emergence risk at forest edges.

## Introduction

Spillover of arthropod-borne viruses (arboviruses) from enzootic foci, facilitated by bridge vectors that feed on both wildlife and humans, can result in isolated human infections or in local epidemics driven by competent urban vectors^1^. Subsequent human-mediated translocations can lead to larger outbreaks with potential global reach^2^. The introduction of pathogens into new geographic regions creates a risk of spillback into sylvatic cycles which, once established, pose a long-term threat to human health through spillover infections^1,2^. All of the arboviruses of greatest public health importance, including yellow fever (YFV), dengue (DENV), Zika (ZIKV) (all *Flaviviridae*: *Orthoflavivirus*), and chikungunya (CHIKV) (*Togaviridae*: *Alphavirus*)^3,4^, originated via spillover from ancestral sylvatic cycles in Africa and Southeast Asia involving monkeys and canopy-dwelling mosquitoes^1,5^ and subsequently established urban transmission cycles in humans sustained by *Aedes* species mosquitoes. YFV was translocated from Africa in the 1700s via the slave trade and established an endemic sylvatic cycle in the Americas involving neotropical monkeys and sylvatic *Haemagogus* and *Sabethes* species mosquitoes^6,7^.

Forest edges, where humans, mosquitoes, and wildlife overlap may serve as a nexus for zoonotic arbovirus exchange^8^. Studies have investigated shifts in mosquito communities^9–12^ and wildlife^13–15^ from interior forest to forest edge, and then to human-modified landscapes. Despite suggestions that mosquito diversity should peak in edge habitats^10^, where urban and sylvatic species overlap, empirical studies have not consistently found highest diversity at edges^9,10,16,17^. However, mosquito species composition and potential routes of spillover and spillback have been shown to change rapidly within a few hundred meters of the edge^17,18^. Several studies have reported higher diversity of wildlife inside forest or at forest edges than in disturbed habitat^13–15^ and have shown that land cover bordering edges may influence wildlife composition^14^. Notably, forest edges influence monkey distributions^19,20^ and risk of interaction with known vectors^18^. Howler monkeys (*Alouatta* spp.), major reservoirs of YFV, are mainly found in the mid to upper forest canopy^20^ and display varying edge associations probably linked to the availability of food^19–21^. Tamarins (*Saguinus* spp.), squirrel monkeys (*Saimiri* spp.), and capuchins (*Sapajus* spp.) are found in the lower canopy and understory, and often in edge habitats^15,20,22^, sometimes venturing into human-modified landscapes^15,23–25^ where they may encounter high densities of dominant vector species, including *Aedes aegypti* and *Ae. albopictus*^10,26^.

Although the likely importance of forest edges for spillover is well recognized, they are often treated as uniform interfaces, regardless of their landscape context. However, forest edges with high habitat contrast, such as those bordering urban land cover, may experience strong edge effects^27^ including a loss of large trees^28^ providing large fruits preferred by larger monkeys^29^, as well as oviposition sites for tree-hole breeding *Haemagogus* and *Sabethes* mosquitoes^30^. Conversely, forest edges with lower habitat contrast, such as those bordering rural or agricultural land cover, may experience weaker edge effects^27^, allowing habitat to remain suitable for forest interior species^31^ including known arbovirus vectors.

Our recent studies of mosquito communities in rainforest fragments bordering Manaus in the Brazilian Amazon^17,18,32,33^ revealed a high relative abundance of *Ae. albopictus* near urban edges^17,18^. The mean Normalized Difference Built-up Index (NDBI), a remote sensing index used to map urbanized areas, was higher in a 100 m radius around sites where this species was present compared to where it was absent when sampling with BG-Sentinel traps^17^, while the opposite trend was observed for *Sabethes* mosquitoes. The presence of *Haemagogus* mosquitoes was not associated with mean NDBI but these were rarely sampled at anthropogenic edges using traps^17^. However, *Haemagogus janthinomys*, a typically canopy-dwelling species and a major neotropical vector of YFV, was frequently encountered at natural edges formed by treefall gaps when sampling at ground level using hand-nets^33^.

In this study, we compared communities of diurnally active, anthropophilic mosquitoes and environmental variables at forest edges bordering urban versus rural land cover in Manaus. We also compared these anthropogenic edges with natural edges formed by treefall gaps as well as with sites in continuous forest. We hypothesized that differences in habitat contrast and environmental variables associated with landscape context would shape differences in mosquito communities. We predicted that diversity would be highest at rural edges owing to overlap of both urban and forest species and that forest edges would alter the vertical stratification of typically canopy-dwelling vectors, including *Haemagogus* and *Sabethes* species. Both changes would have significant implications for the risk of spillover and spillback.

## Methods

### Study area

The study was carried out at the Adolpho Ducke forest reserve^34^ (Ducke) near Manaus, a city of more than two million people situated at the confluence of the Negro and Solimões rivers in the Brazilian Amazon (Figure 1). Ducke is 100 km^2^ of *terra firme* rainforest home to multiple mammal species including six species of monkeys which differ in their ecology and behavior^35^. The reserve forms an abrupt border with the city along its southwestern edge, where people live in close contact with wildlife and sylvatic mosquitoes^17^. The remaining border of Ducke abuts rural areas, where edges may be similarly abrupt, or where the transition between primary and secondary vegetation and nearby habitations is more gradual. Rural areas are mostly characterized by villages and other smallholdings where produce includes cupuaçu (*Theobroma grandiflorum*), açaí (*Euterpe oleracea*), and andiroba (*Carapa guianensis*)^36^, and animals such as dogs and chickens are commonly kept. Anecdotally, in both urban and rural areas, residents occasionally enter the forest for gathering fruit, hunting, or bathing in streams^17^. DENV, ZIKV, and CHIKV circulate in urban cycles in Manaus^37^, while YFV^38^ and Mayaro virus (MAYV, *Togaviridae*: *Alphavirus*)^39^ circulate in nearby forests. Mosquito abundance is highest during the rainy season, which usually lasts from November until May, and decreases over the drier period from June until October^34^.

**Figure 1.**
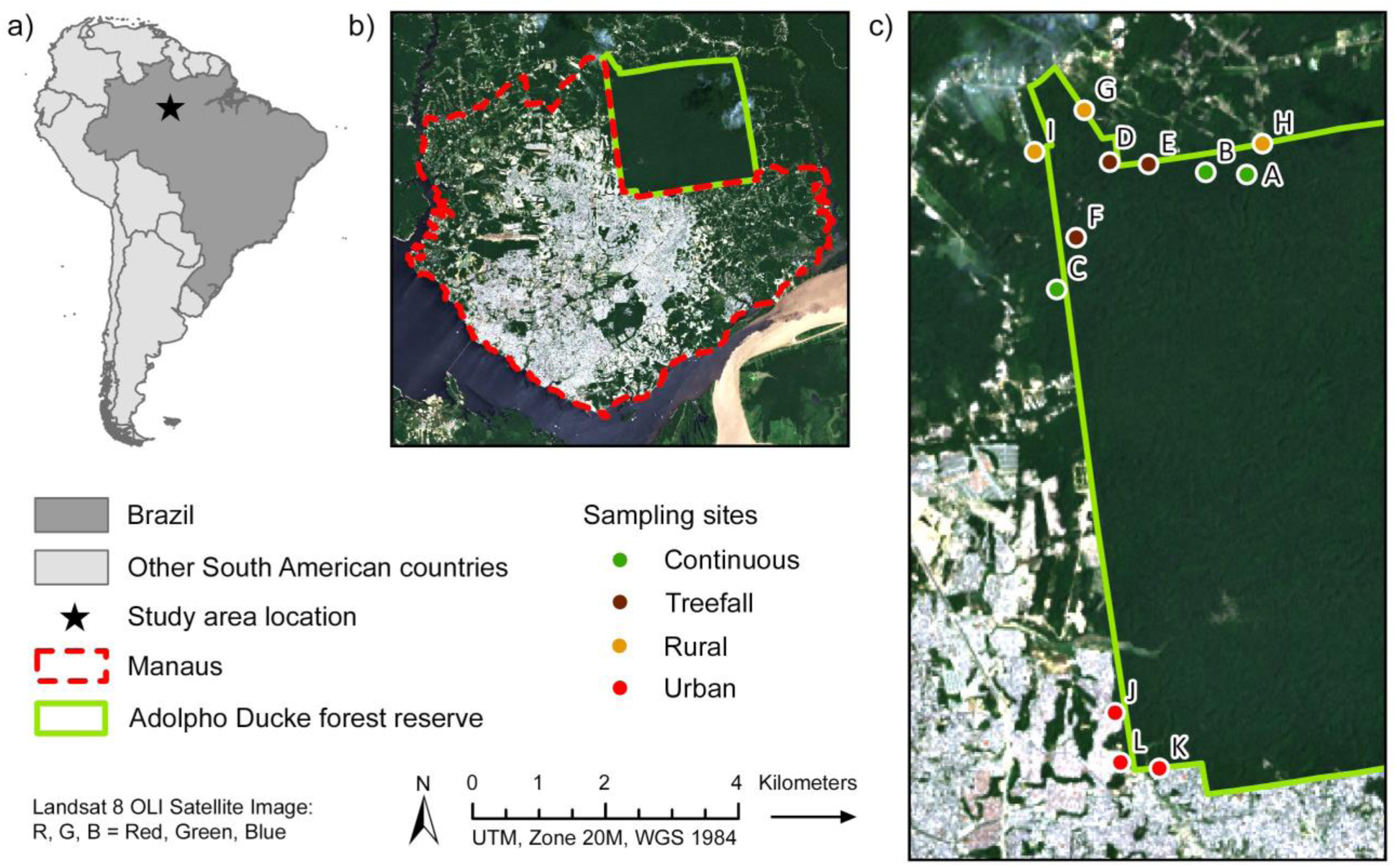
Reference map of the study area based on Landsat 8 Operational Land Imager (OLI) surface reflectance imagery obtained from the USGS Earth Explorer data portal^40^. Panels show a) the location of Manaus in Brazil, b) the Adolpho Ducke forest reserve on the edge of Manaus, and c) the location of sampling sites in each category. The Manaus polygon was derived from a GISMAPS neighborhoods shapefile^41^, while the Ducke reserve boundary was provided by the Instituto Nacional de Pesquisas da Amazônia (INPA). The map was created using ArcGIS Desktop 10.7.1 (ESRI, Redlands, California).

### Forest edge type and site selection

Three biologically independent sampling sites, located by direct surveillance, were situated in each of three types of forest edge: edges bordering urban land cover, edges bordering rural land cover, and internal edges formed by natural treefall gaps. A further three sites situated in areas of continuous forest served as controls (Supplementary Figure 1). Thus, the study comprised 12 sites in total. Urban and rural edge sites were established where deemed safe, accessible, and in agreement with local communities. Urban edge sites were close to built-up residential areas within 10 m of the nearest house. Rural edge sites were in more sparsely populated village and agricultural areas at least 40 m from the nearest house. Treefall gap and continuous forest sites were 500 m from the forest edge and were different from those sampled in previous studies^32,33^. Treefall gap sites were characterized by an opening in the canopy caused by one or more fallen trees (> 10 m in height) allowing a moderate to significant amount of sunlight to reach the forest floor. Continuous forest sites were beneath intact canopy allowing very little sunlight to reach the forest floor. The mean NDBI value in a 100 m buffer surrounding each sampling site was calculated as previously described^17^. All sampling sites were situated at least 500 m apart to minimize spatial autocorrelation.

### Sampling platforms

A five-meter-high timber platform was constructed between two nearby trees within two meters of the forest edge at edge sites, or beneath intact canopy at continuous forest sites. We chose this height as we previously found significant changes in mosquito communities between ground level (0 m) and 5 m when sampling at a treefall gap inside the same forest^33^. We also saw significant breakpoints in temperature and relative humidity between these heights when sampling beneath the forest canopy^32^.

### Mosquito collections

Mosquitoes were sampled using hand-nets over 12 months beginning in the early dry season on 6 July 2021 and ending on 30 June 2022. When sampling a site, two collectors worked simultaneously, one at ground level and one on the platform, to collect all approaching mosquitoes between 10:00 and 15:00, when the vector species of primary interest tend to be active^42,43^. These were aspirated and separated into 50 mL Falcon tubes at 30-minute intervals. Tubes containing live mosquitoes were placed in a Styrofoam box in the shade to prevent desiccation until they were transferred to a −80°C freezer at the Fundação de Medicina Tropical Doutor Heitor Vieira Dourado (FMT-HVD) at the end of each day. The height at which a collector worked was generally alternated daily to minimize effects of variation in collector attractiveness. We aimed to sample one site per day, three days per week, rotating between sites in a 12 x 12 Latin square design.

### Microclimate, weather, and rainfall

Environmental variables were recorded to investigate their associations with edge type and/or the abundance and occurrence of key mosquito taxa. Hygrochron iButton data loggers (Maxim Integrated, San Jose, California) were used to record temperature (°C) and relative humidity (%) at each height at 15 and 45 minutes past each hour (i.e., the midpoint of each 30-minute interval). iButtons were placed in nylon mesh bags; one was hung from vegetation close to the collector at ground level while the other was hung above the platform. Data were additionally used to calculate the daily minimum, maximum, mean, and range of both temperature and relative humidity variables. Weather was manually recorded at 30- minute intervals in dry conditions as 1 = clear skies, 2 = scattered cloud, 3 = overcast, and in wet conditions as 4 = light rain, or 5 = heavy rain. Collections were suspended in stormy conditions. The daily mean weather was then calculated, with values closer to 1 indicating favorable weather and values closer to 5 indicating inclement weather. Precipitation data, obtained from an automated meteorological station^44^ (INMET code: Manaus-A101, OMM: 81730, 3.103682° S, 60.015461° W), were used to calculate 7-day cumulative rainfall lagged at 1, 2, 3, and 4 weeks prior to each sampling day. In this study, we defined the rainy season as November until April when cumulative monthly rainfall consistently exceeded 250 mm^44^; this is a slightly shorter period that the standard regional rainy season of November to May.

### Mosquito identifications

Mosquitoes were placed on a chill table (BioQuip, Rancho Dominguez, California, USA) and morphologically identified by Mr. Nelson Fé, who was unaware of the site of origin of the specimens, using a stereomicroscope and taxonomic keys as previously described^32^. Genus and species names and respective abbreviations follow Wilkerson et al.^45^. Samples were stored at -80°C for future arbovirus screening.

### Statistical analysis

Statistical analyses were performed with JMP 17^46^ unless stated. To investigate how microclimate varied across forest edges, by season, and by height of collection, Spearman’s rank correlation was first used to identify significantly associated variables, of which, mean temperature and mean relative humidity were chosen for further analysis. To compare microclimate between two groups: forest edges (N = 9) vs. continuous forest (N = 3), rainy season vs. dry season, and 0 m vs. 5 m, a two-tailed t-test for normally distributed data and a Wilcoxon Rank Sum test for non-normal data were used. To compare microclimate across the three edge habitats for both combined and height specific data, a one-way ANOVA (normal data) and a Kruskal-Wallis test (non-normal data) were used.

Measurements of mosquito community similarity and diversity were based on specimens identified to the rank of species. For these analyses, data were grouped by 1) edge type, 2) edge type and height, or 3) edge type and season (at 0 m and 5 m separately). The Morisita overlap index, based on species count data, was calculated using the PAST version 4.14 software package^47^ to compare mosquito community composition between sampling sites for each edge type, and for data grouped as described above. To compare the similarity of communities based on relative species abundance data, Spearman’s rank correlation was first used to identify highly significantly correlated species. Where significance was P ≤ 0.01, the least abundant of the two species was excluded. The resulting datasets were then used for principal components analysis followed by hierarchical clustering of the principal components.

To estimate species richness and examine whether sampling was adequate to capture total richness, iNEXT^48^ (R version 4.2.2) was used to generate rarefaction curves by edge type for each height and season sampled. Species evenness was calculated for each site as Shannon-Wiener diversity index (*H’*) divided by the natural logarithm of species richness^10^. Since data were normally distributed, a one-way ANOVA followed by a Tukey HSD post-hoc test was used to compare evenness by edge type. A two-way ANOVA was used to compare evenness by edge type and height, and edge type and season, followed by a post-hoc least-squares means Tukey HSD test to compare multiple means (i.e., edge type) or a least-squares means t-test to compare two means (i.e., season). A one-way ANOVA was used for simple effects tests when interaction effects were detected.

A Wilcoxon Rank Sum test (non-normal data) was used to compare differences in mosquito abundance between rainy season vs. dry season at each edge type for each of the eight most abundant species sampled overall. Data from both heights were combined, and comparisons with fewer than 60 mosquitoes (an arbitrary cutoff to reduce random sampling effects) were excluded.

Nominal logistic regression was used to test associations between the occurrence (presence/absence) of each of the eight most abundant species with environmental variables chosen based on our field observations and previous work^17,32^. These were: edge type, mean weather, mean temperature, mean relative humidity, height, and 7-day cumulative rainfall lagged at 1, 2, 3, and 4 weeks. Variables were removed sequentially from the model until all remaining variables contributed significantly or only one variable was left. Due to the high number of variables, we chose an alpha value of 0.01 as highly significant for this analysis, while 0.01 > P < 0.1 was considered marginally significant, and tested associations in each season separately. We supplemented this analysis by testing the effect of edge type and height on species occurrence using a generalized linear model with a normal distribution and an identity link function, based on the % positive sampling days at ground level and platform sites (thus N = 6 in total) for each edge type. If the interaction effect was significant, we conducted a Kruskal-Wallis Rank Sum test to analyze simple effects of edge type for each height separately. We additionally used a standard least squares analysis to test rainy season associations between environmental variables and abundance of the eight species. For this analysis, sampling sites were nested into edge type, which was not included as a variable, although edge types with significantly lower abundance were excluded. Variables included in the model were otherwise the same as described above. We did not test dry season associations with species abundance due to data being heavily zero inflated.

Prompted by our field observations, we used contingency tables and a Pearson’s chi-square test for large frequencies to further explore relationships between *Sabethes* mosquitoes grouped at subgenus level (*Sabethes* or *Sabethoides*), height, and edge type using 30-minute occurrence data^49^.

### Ethics and permits

Mosquito collections at the Ducke reserve were approved by local environmental authorities (SISBIO license 57003-6) and the study did not involve endangered or protected species. When collecting with hand-nets, skin was not deliberately exposed to attract mosquitoes and mosquito landing was not permitted. Collectors are listed among the co-authors and were fully aware of the nature of the research. They wore trousers, a long-sleeved shirt and/or repellent to minimize the risk of being bitten and had been vaccinated against yellow fever.

## Results

### Mean NDBI varied most between urban edge sites

The mean NDBI values within a 100 m buffer surrounding each sampling site ranged from -0.223 to -0.532 at urban edges, -0.582 to -0.629 at rural edges, -0.666 to -0.700 at treefall gaps, and -0.696 to -0.728 in continuous forest. There was a substantial amount of forest cover within the 100 m buffer at all edge sites. As a result, NDBI values were relatively low across all categories.

### Forest edges, irrespective of landscape context, were hotter and drier than continuous forest

The fluctuation in temperature and humidity across the daily sampling period (10:00 – 15:00) differed between forest edges and continuous forest and differed among forest edges depending upon landscape context and season (Figure 2). Mean temperature and relative humidity were significantly correlated with all other microclimate variables (Spearman’s rank correlation, P < 0.0001). Both variables remained stable throughout the sampling hours in continuous forest but fluctuated considerably at treefall gaps, which were hottest and driest during the early afternoon hours. The magnitude of diel fluctuations in microclimate at rural and urban edges was intermediate between continuous forest and treefall gaps. As expected, mean temperature at forest edges was significantly higher than in continuous forest (two-tailed unequal variances t-test, DF = 169.2, *t* = 3.87, P = 0.0002) and was significantly higher in the dry season than in the rainy season (DF = 277.7, *t* = -5.57, P < 0.0001). The inverse was true for mean relative humidity (Wilcoxon Rank Sum, DF = 1, χ^2^ = 14.5, P = 0.0001 and DF = 1, χ^2^ = 47.3, P < 0.0001, respectively). When data were combined for both heights, neither mean temperature (one-way ANOVA, DF = 2, F = 1.42, P = 0.24) nor mean relative humidity (Kruskal-Wallis, DF = 2, χ^2^ = 5.92, P = 0.052) differed across the three edge habitats. The same was true when these variables were compared at each height separately (P > 0.1 for all comparisons). In addition, conditions were marginally hotter (two-tailed equal variances t-test, DF = 217, *t* = 1.93, P = 0.06) and were drier (Wilcoxon Rank Sum, DF = 1, χ^2^ = 11.1, P = 0.0009) 5 m above the ground than at ground level when data were combined across the three edge habitats.

**Figure 2.**
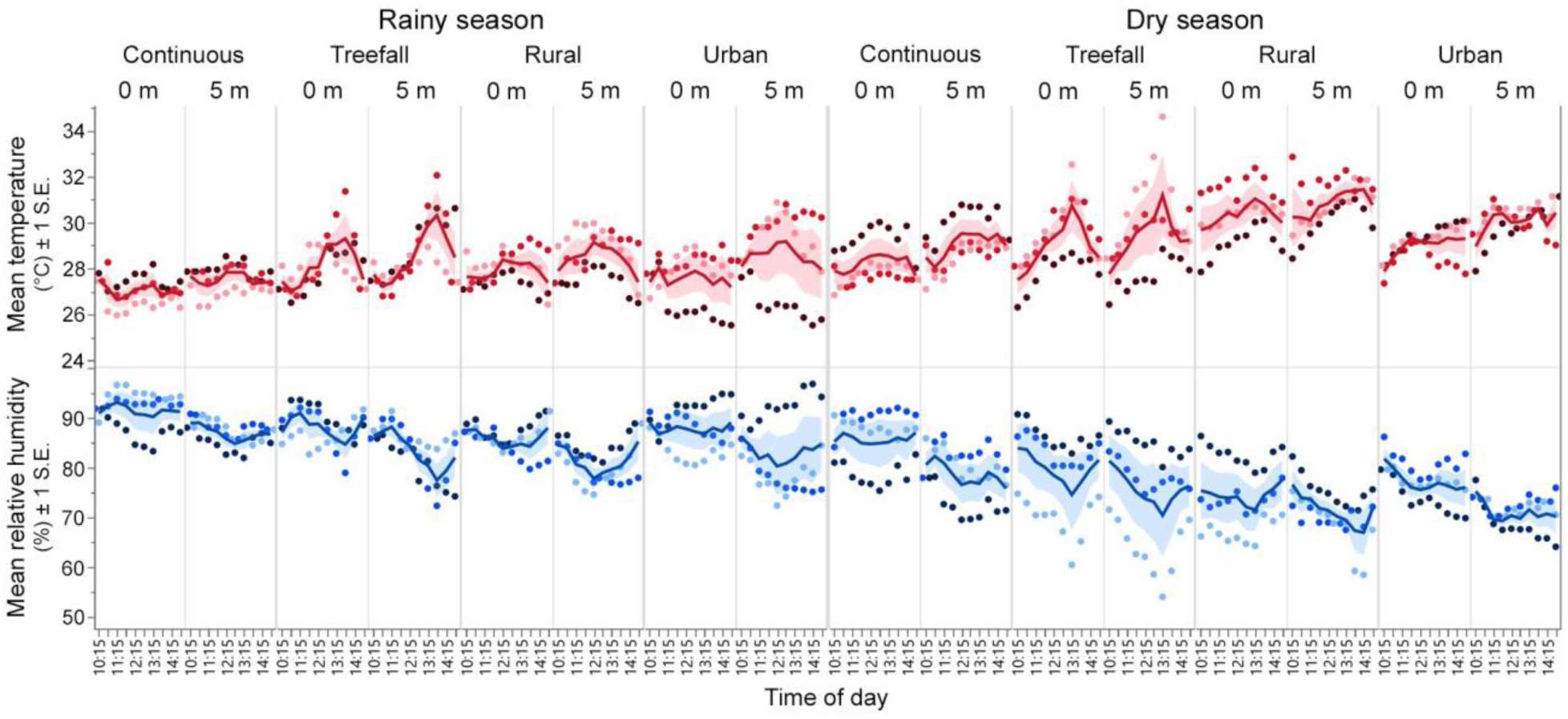
Variation in microclimate by season, edge type, and height during the daily sampling period. Colored dots represent the mean temperature (°C) and mean relative humidity (%) for the three sites sampled in each edge type at each designated timepoint. Solid lines represent the mean of the three site values and shaded areas show ± 1 standard error (S.E.).

### Overview of mosquito collections

Mosquito sampling was conducted over 71 rainy season days and 75 dry season days. Sampling effort was relatively evenly distributed across categories, with 36 days spent in continuous forest, 37 days at treefall gaps, 35 days at rural edges, and 38 days at urban edges. Collections yielded 4,425 adult mosquitoes (97.5% female, 13 genera, and 69 identified species) including 1,503 in continuous forest, 1,682 at treefall gaps, 629 at rural edges, and 611 at urban edges (Figure 3, Dataset^49^). Of these, 2,524 were sampled at ground level and 1,901 at 5 m, while 2,854 were sampled during the rainy season and 1,571 during the dry season. The most abundant genera were *Haemagogus* (30.2 %), *Psorophora* (21.6 %), *Sabethes* (16.2 %), *Limatus* (12.3 %), and *Wyeomyia* (9.6 %), while *Aedes* mosquitoes formed 5.1% of the total catch. The most abundant species were *Hg. janthinomys* (27.0 %), *Ps*. *amazonica* (19.2 %), *Li. durhamii* (7.5 %), *Sa. chloropterus* (5.8 %), *Wy. aporonoma* (3.2 %), *Ae*. *albopictus* (3.1 %), *Sa. cyaneus* (2.6%), and *Li. pseudomethysticus* (2.5%).

**Figure 3.**
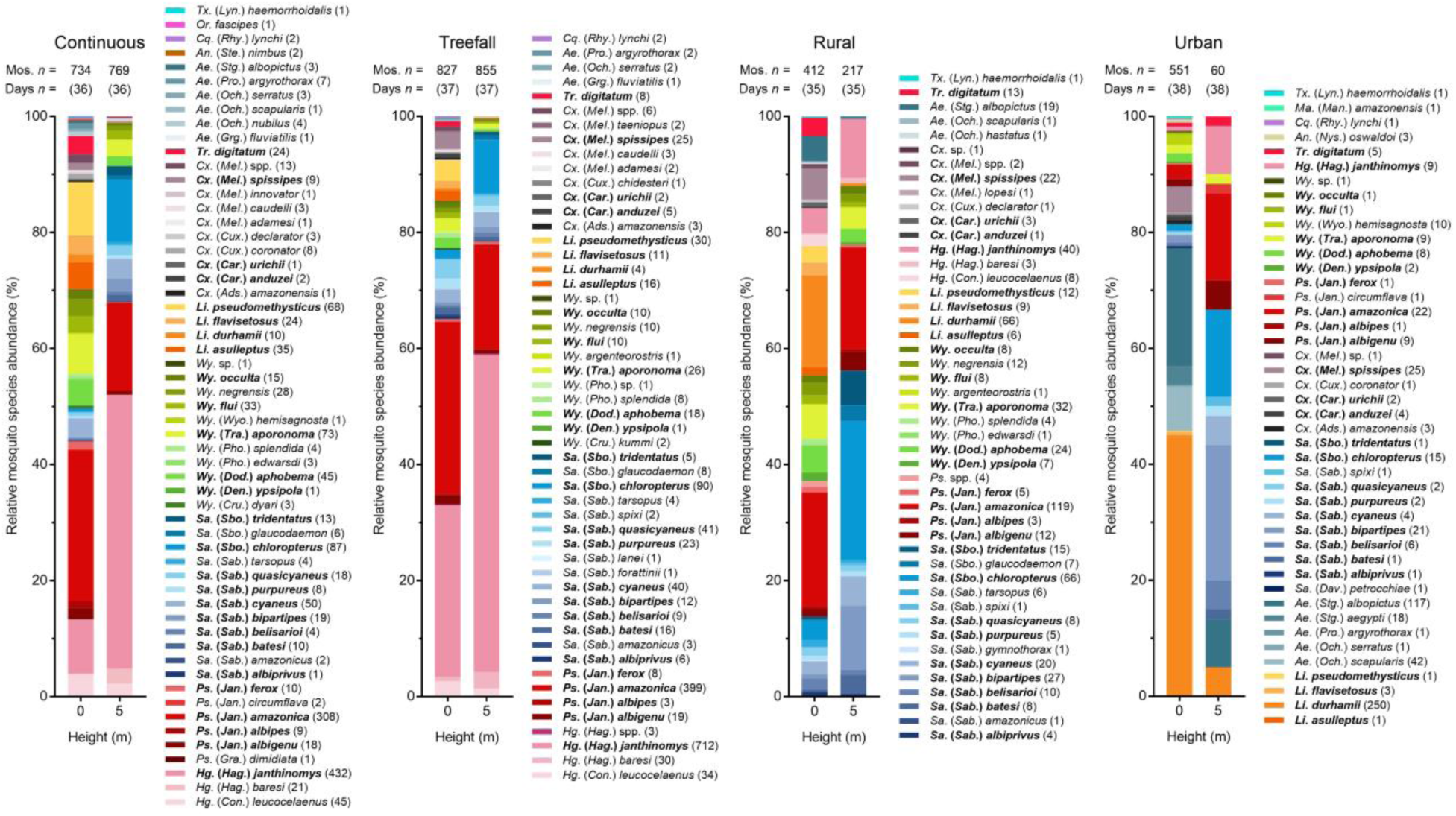
Relative mosquito species abundance by edge type and height of collection. Stacks ordered by genus abundance and then alphabetically by subgenus and species. Number of mosquitoes (Mos. *n* =) and number of sampling days (Days *n* =) in each edge type and at each height shown above bar. Number of individuals per taxon included in parentheses next to corresponding name; sp. = single species, spp. = potentially multiple species. Abbreviated names are given in full in the Dataset^49^. Species collected across all four categories highlighted in bold text.

### Landscape context and height together shaped the mosquito community composition in different edge habitats

*Haemagogus janthinomys* and *Ps. amazonica* dominated collections in continuous forest and at treefall gaps, where *Sabethes* species, including *Sa. chloropterus*, were also common (Figure 3). These taxa were particularly abundant at 5 m, while at ground level, *Wyeomyia* and *Limatus* also formed a high proportion of the catch. The relative abundance of *Wyeomyia* and *Limatus* at ground level was more than twice as high in continuous forest than at treefall gaps. The reverse was true for *Sabethes* species, while the ground level relative abundance of *Hg. janthinomys* was 3.5 times higher at treefall gaps than in continuous forest. At rural edges, the relative abundance of mosquitoes at ground level was more evenly distributed among genera, with *Sabethes*, *Psorophora*, *Wyeomyia*, and *Limatus* being well represented. At urban edges, *Limatus and Aedes* dominated collections. *Psorophora amazonica*, *Li. durhamii*, and *Ae. albopictus* were the dominant species within their genera at rural and urban edges. In these settings, *Sabethes* species formed more than 50% of the mosquitoes sampled on platforms.

The Morisita index revealed high overlap between sites in continuous forest and at treefall gaps, but lower overlap between sites at rural edges and at urban edges (Supplementary Table 1). Based on species collected at both ground level and 5 m above the ground combined (Table 1), mosquito communities in continuous forest were similar to those at treefall gaps but differed greatly from communities at the urban edge. Communities at rural edges were moderately similar to all other edge types. When this analysis was broken down by height (Table 2), ground level community composition followed the pattern described above. At 5 m above the ground, however, continuous forest and treefall gap communities were almost indistinguishable; rural edges and urban edges showed substantial overlap with each other, and both showed moderate overlap with interior forest communities. There was little change between rainy and dry season in mosquito community composition at ground level or 5 m, although urban edges were excluded from the latter comparison due to small sample size (Supplementary Table 1).

**Table 1.** Morisita overlap index for comparisons by edge type. Cont. = Continuous forest.

|  | Cont. | Treefall | Rural | Urban |
| --- | --- | --- | --- | --- |
| Cont. | 1 |  |  |  |
| Treefall | 0.939 | 1 |  |  |
| Rural | 0.686 | 0.524 | 1 |  |
| Urban | 0.107 | 0.083 | 0.445 | 1 |

**Table 2.**
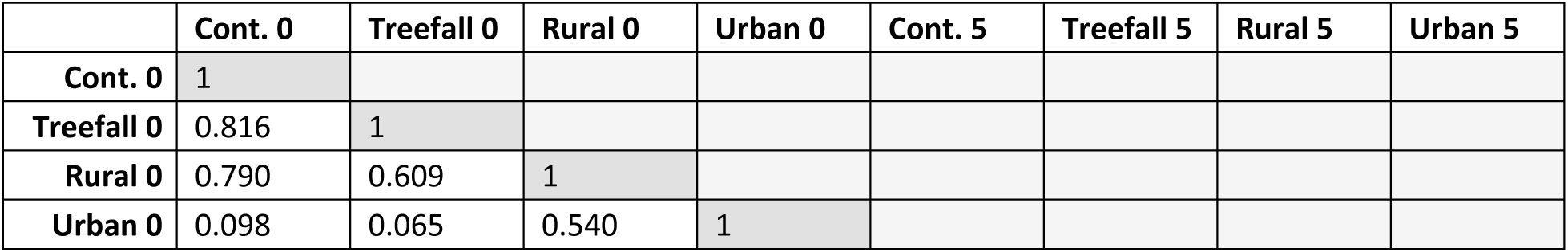
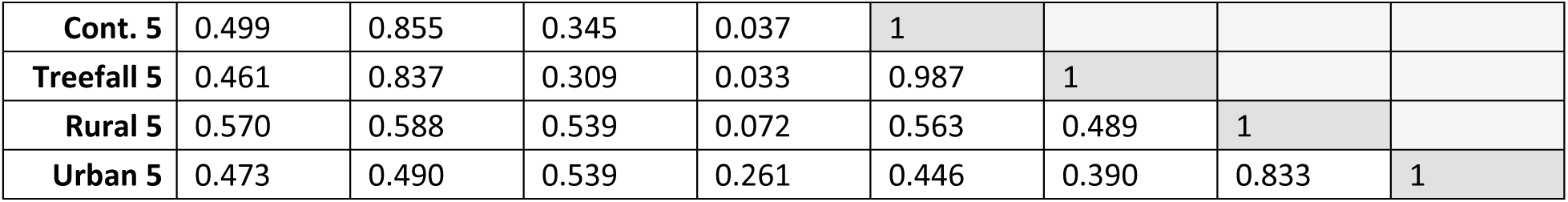
Morisita overlap index for comparisons by edge type and height. Cont. = Continuous forest, 0 = 0 m, 5 = 5 m.

To conduct a principal components analysis of relative species abundance at each edge type for both heights combined, we first checked pairwise correlations of each species and removed one species from each pair that was highly significantly (P ≤ 0.01) correlated. Principal component (PC)1 and PC2 captured 40.7% and 38.1% of the variation in data, respectively (Figure 4). PC1 represented the relative abundance of *Hg. janthinomys*, *Hg. leucocelaenus*, *Sa. bipartipes*, and *Sa. belisarioi*, along with rarer species including *Cx. caudelli* and *Cx. adamesi*, while PC2 represented the relative abundance of a group of *Sabethes* species containing *Sa. chloropterus*, *Sa. batesi*, and *Sa. albiprivus*, along with *Wy. argenteorostris* and *Ae. hastatus* (Supplementary Table 2). In a principal components analysis of relative species abundance at each edge type and height, PC1, PC2, and PC3 captured 23.1%, 22.4%, and 16.8% of the variation in data, respectively. PC1 represented the relative abundance of several *Sabethes* species including *Sa. cyaneus*, *Sa. purpureus*, and *Sa. chloropterus*, along with *Wy. hemisagnosta*, *Ae. aegypti*, and *Ae. scapularis*. PC2 represented the relative abundance of *Wy. aporonoma*, *Ps. amazonica*, *Hg. leucocelaenus*, and *Tr. digitatum* among other predominantly rarer species, while PC3 represented the relative abundance of *Wy. ypsipola* and *Hg. janthinomys* among the highest loading species. We did not perform principal components analysis by season since we detected little seasonal change in mosquito community composition using the Morisita index.

**Figure 4.**
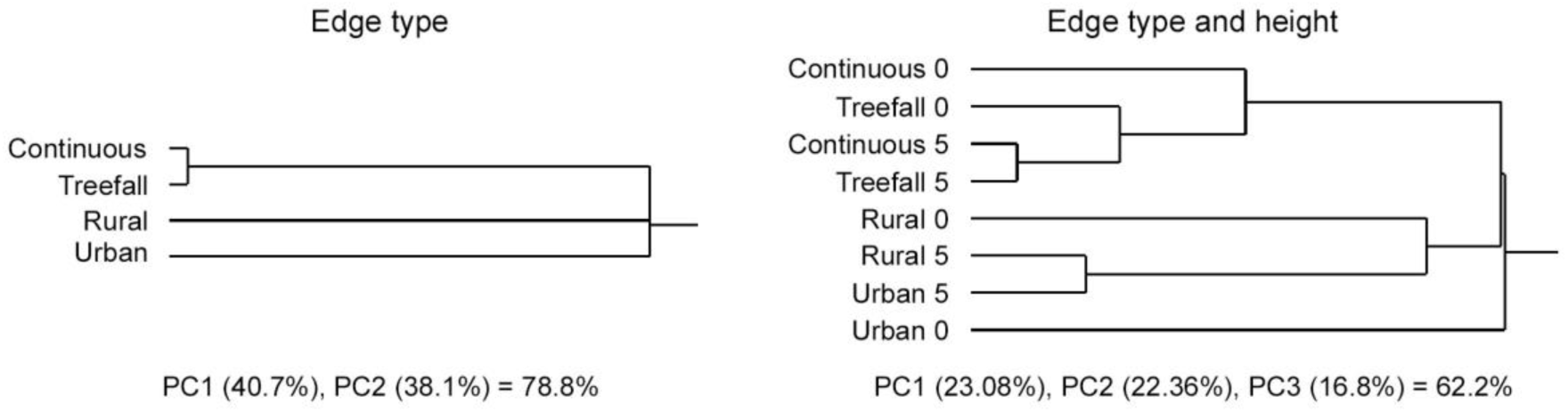
Mosquito community structure. Dendrograms show hierarchical clustering of principal component PC1 and PC2 from the analysis of relative species abundance by edge type, and PC1, PC2, and PC3 from the analysis by edge type and height (0 = 0 m, 5 = 5 m).

Results of hierarchical clustering generally agreed with Morisita comparisons. Hierarchical clustering of PC1 and PC2 by edge type showed that continuous forest and treefall gap communities were very similar, but differed considerably from rural edge communities, which in turn differed from urban edge communities (Figure 4). For the edge type and height comparison, hierarchical clustering of PC1, PC2, and PC3 showed that continuous forest and treefall gap communities were more similar to each other than to the cluster of rural edge and 5 m urban edge communities, while 0 m urban edge communities were distinct from both clusters. Continuous forest and treefall gap communities 5 m above the ground were very similar and more closely resembled ground level treefall gap communities than ground level continuous forest communities.

### Landscape context shaped species diversity at forest edges

A total of 55 identified species were sampled in continuous forest, 48 at treefall gaps, 44 at rural edges, and 42 at urban edges. Urban edges exhibited the greatest variation in species richness, ranging from 14 to 33 species per site. Rarefaction and extrapolation curves showed little difference in estimated species richness across edge types, although sampling was insufficient to capture total species richness at rural and urban edges (Supplementary Figure 2). Mean species evenness, on the other hand, differed significantly across edge types (one-way ANOVA, DF = 3, F = 9.46, P = 0.005), with rural edges having significantly higher evenness than urban edges and treefall gaps (Supplementary Table 3). When data were analyzed separately at each height, estimated species richness followed the same pattern but was higher at 0 m than at 5 m overall (Supplementary Figure 2). Sample coverage at rural and urban edges was also higher at 0 m than at 5 m. There was a significant interaction between edge type and height for species evenness (two-way ANOVA, DF = 3, F = 21.8, P < 0.0001). Simple effects tests showed that evenness differed across edge types at both 0 m (one-way ANOVA, DF = 3, F = 9.78, P < 0.005) and 5 m (DF = 3, F = 33.5, P < 0.0001). At 0 m, evenness was significantly higher at rural edges than at treefall gaps and urban edges (Table 3), and higher in continuous forest than at urban edges. At 5 m, evenness was higher at both rural and urban edges than at both treefall gaps and in continuous forest.

**Table 3.** Species richness, diversity, and evenness by edge type and height. Arithmetic means (± 1 standard error) calculated per sampling site (N = 3).

| Edge type | N = | Richness | | Diversity ($H'$) | | Evenness | |
| --- | --- | --- | --- | --- | --- | --- | --- |
|  |  | 0 m | 5 m | 0 m | 5 m | 0 m | 5 m |
| Continuous | 3 | 33.0 (1.15) | 21.0 (0.58) | 2.76 (0.01) | 1.95 (0.14) | 0.79 (0.01) | 0.64 (0.05) |
| Treefall | 3 | 31.7 (2.40) | 21.3 (2.19) | 2.29 (0.18) | 1.65 (0.11) | 0.66 (0.04) | 0.54 (0.02) |
| Rural | 3 | 30.0 (0.58) | 15.3 (4.67) | 2.81 (0.09) | 2.14 (0.27) | 0.83 (0.03) | 0.83 (0.03) |
| Urban | 3 | 19.7 (6.17) | 8.70 (1.33)* | 1.75 (0.05) | 1.93 (0.13)* | 0.62 (0.04) | 0.91 (0.02)* |

$H'$ = Shannon-Wiener diversity index.
\*Small sample size at 5 m urban edge (60 mosquitoes).

Estimated species richness followed the same general pattern described above during the rainy season (Figure 5). During the dry season, rarefaction and extrapolation curves diverged at 0 m, where richness was highest at rural edges and in continuous forest. Species richness was higher at treefall gaps than in continuous forest when sampling at 5 m, while sample coverage was low at rural edges. At 0 m, there was a significant effect of edge type (two-way ANOVA, DF = 3, F = 11.8, P = 0.0003) and season (DF = 1, F = 7.49, P = 0.015) on species evenness (Supplementary Table 3), but no interaction effect between these variables (DF = 3, F = 0.12, P = 0.9). Evenness at rural edges and in continuous forest was again higher than at treefall gaps and urban edges and was slightly but significantly lower in the rainy season than in the dry season. At 5 m, there was a significant effect of edge type (two-way ANOVA, DF = 2, F = 21.6, P = 0.0001), but no effect of season (DF = 1, F = 3.98, P = 0.07) or interaction between these variables (DF = 2, F = 2.12, P = 0.2). At this height, evenness was higher at rural edges than in continuous forest and at treefall gaps.

**Figure 5.**
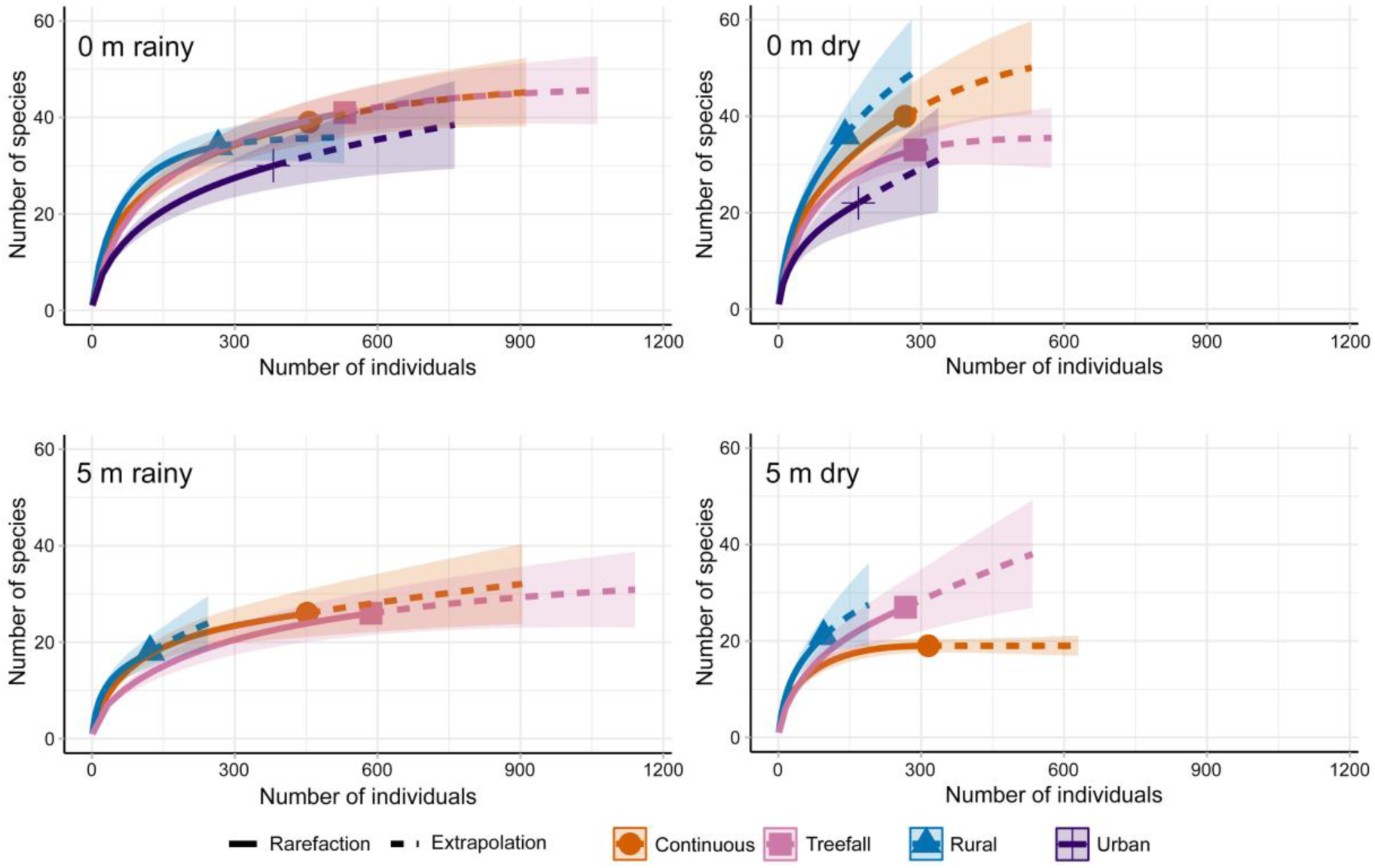
Species richness rarefaction and extrapolation curves. Panels show richness for data grouped by edge type (based on N = 3 biologically independent sampling sites per edge type) for each height and season. Shaded areas surrounding rarefaction and extrapolation lines represent 95% confidence intervals. Urban edges were excluded at 5 m due to small sample size.

### Mosquito abundance decreased but key vector species persisted through dry season months

Comparisons of rainy season vs. dry season abundance at each edge type for the eight most abundant species revealed significant differences only for *Hg. janthinomys* and *Sa. chloropterus* at treefall gaps (Wilcoxon Rank Sum, DF = 1, χ^2^ = 12.5, P = 0.0004 and DF = 1, χ^2^ = 3.84, P = 0.0499, respectively) and *Li. durhamii* at rural edges (DF = 1, χ^2^ = 9.43, P = 0.002). In all cases, the median number of mosquitoes was significantly higher during the rainy season (Figure 6, Supplementary Table 4). The contrast between seasons was most pronounced for *Hg. janthinomys* sampled at treefall gaps, although *Ps. amazonica* exhibited a tendency towards higher rainy season abundance in continuous forest, at treefall gaps, and at rural edges. While the overall mosquito abundance tended to be higher during the rainy season, several species, including *Hg. janthinomys* and *Ae. albopictus*, maintained appreciable numbers throughout the dry season. There was little difference between the number of *Sa. chloropterus* sampled in rainy season vs. dry season months in continuous forest (DF = 1, χ^2^ = 0.04, P = 0.83) and at rural edges (DF = 1, χ^2^ = 0.7, P = 0.4). Even at treefall gaps, where the difference was significant, *Sa. chloropterus* still persisted in relatively high numbers.

**Figure 6.**
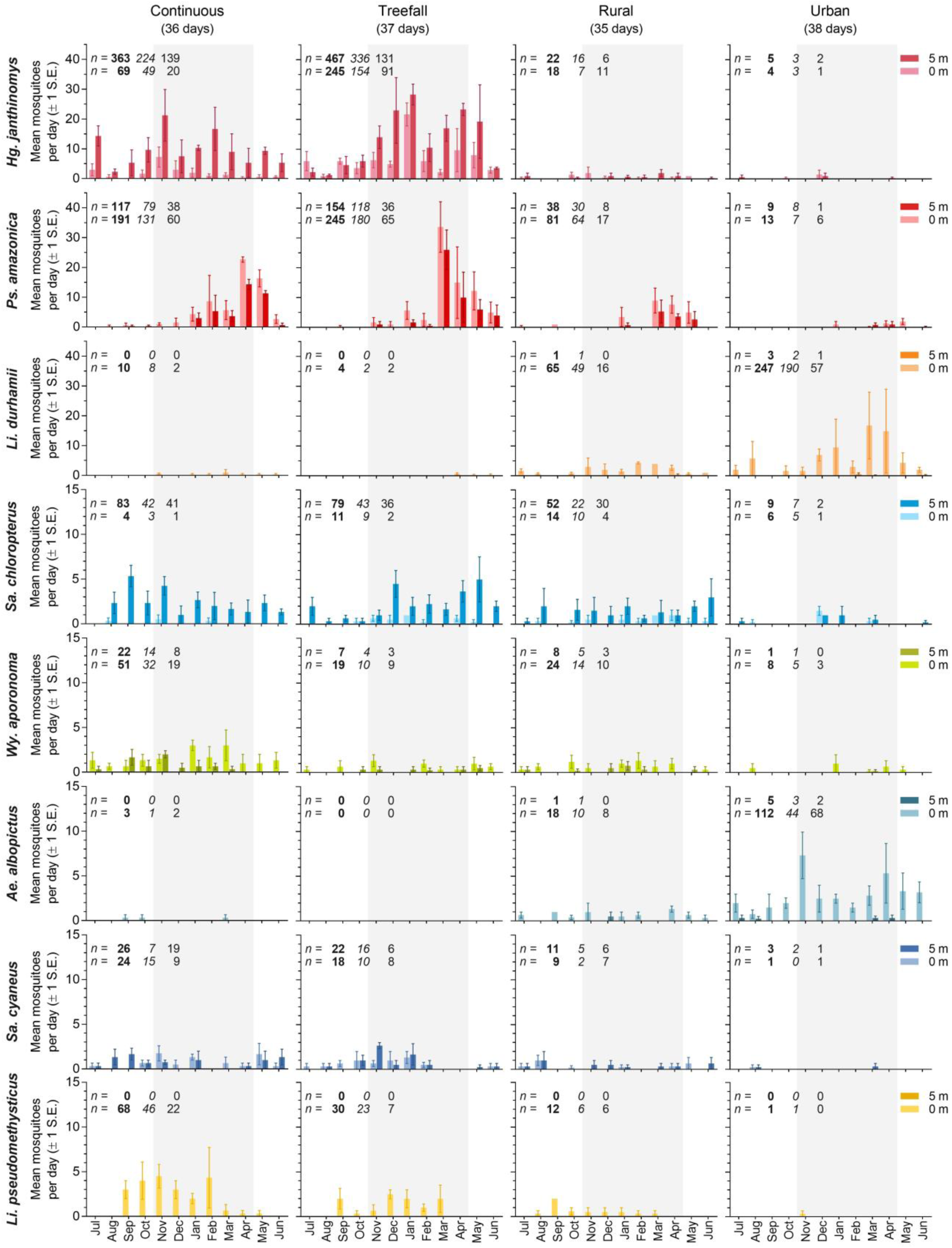
Mean number of mosquitoes sampled per day ± 1 standard error (S.E.) by edge type, height, and month of collection for the eight most abundant species. Collections were made from July 2021 – June 2022. Gray shaded areas show rainy season months (November 2021 – April 2022). Note that species 1 – 3 are plotted on a different y-axis to species 4 – 8. *n* = shows the total (bold font), rainy season (italic font) and dry season (regular font) number of mosquitoes sampled at 0 m (bottom row) and 5 m (top row) for each species.

### Occurrence and abundance of key vectors showed species-specific associations with edge type and other environmental variables

Nominal logistic regression (Table 4) showed that edge type effects on occurrence of the same eight species were almost universal but partitioning by edge differed among species. During the rainy season, sylvatic species (all but *Li. durhamii* and *Ae. albopictus*) were prevalent in continuous forest and at treefall gaps, while *Hg. janthinomys*, *Ps. amazonica*, *Sa. chloropterus*, *and Wy. aporonoma* were also common at rural edges (Supplementary Table 5). Of the urban species, *Ae. albopictus* was more prevalent at urban edges than at rural edges, *Li. durhamii* occurred evenly between the two, but neither were common inside the forest. These patterns held true during the dry season, albeit at slightly lower levels for several species. Notably, there was little difference in occurrence across edge types between rainy and dry seasons for *Hg. janthinomys, Sa. cyaneus*, and *Ae. albopictus*.

**Table 4.** Nominal logistic regression (P value and (χ^2^)) testing associations between occurrence of the eight most abundant species overall and environmental variables in rainy and dry seasons. Blue and red shaded cells indicate significant positive or negative associations, respectively. P values in bold font represent highly significant results (P < 0.01). N = 3 biologically independent sampling sites per edge type.

|  |  |  |  |  |  | 7-day cumulative rainfall lag |  |  |  |
| --- | --- | --- | --- | --- | --- | --- | --- | --- | --- |
| Species [*n* =]* | Edge type | Height<sup>†</sup> | Mean temp | Mean RH | Mean weather<sup>‡</sup> | 1 wk | 2 wk | 3 wk | 4 wk |
| RAINY SEASON |  |  |  |  |  |  |  |  |  |
| *Hg. janthinomys* [78] | <0.0001 (97.0) | 0.03 (4.6) | 0.03 (4.6) |  |  | 0.07 (3.3) |  |  |  |
| *Ps. amazonica* [64] | 0.001 (16.2) |  | 0.003 (8.8) |  |  |  |  | 0.0008 (11.2) |  |
| *Li. durhamii* [37] | <0.0001 (30.4) | <0.0001 (39.9) | 0.04 (4.1) | 0.04 (4.4) |  | 0.005 (7.7) |  |  |  |
| *Sa. chloropterus* [66] | 0.0003 (18.6) | <0.0001 (17.4) |  |  | 0.002 (9.7) |  |  |  |  |
| *Wy. aporonoma* [48] | 0.0008 (16.6) | 0.02 (5.3) |  |  | 0.048 (3.9) |  |  |  |  |
| *Ae. albopictus* [29] | <0.0001 (59.1) | <0.0001 (31.9) |  |  |  |  |  |  |  |
| *Sa. cyaneus* [36] | 0.002 (15.0) |  |  |  | 0.005 (8.0) | 0.05 (3.9) |  |  |  |
| *Li. pseudomethysticus* [29] | <0.0001 (23.9) | <0.0001 (57.9) |  |  | 0.02 (5.4) |  | 0.01 (6.2) |  |  |
| DRY SEASON |  |  |  |  |  |  |  |  |  |
| *Hg. janthinomys* [74] | <0.0001 (89.8) | 0.06 (3.5) |  |  |  |  |  |  |  |
| *Ps. amazonica*<br>[34] | **<0.0001**<br>(23.1) | 0.05 (4.0) |  |  | **0.0003**<br>(12.9) |  | **0.002**<br>(9.3) | 0.01 (6.1) | **<0.0001**<br>(19.7) |
| *Li. durhamii*<br>[28] | **<0.0001**<br>(29.7) | **<0.0001**<br>(34.7) |  | 0.02 (5.9) |  |  |  |  |  |
| *Sa. chloropterus*<br>[45] | **0.002**<br>(14.4) | **<0.0001**<br>(47.7) |  | **<0.0001**<br>(15.7) | **0.002**<br>(9.4) |  |  |  | 0.08 (3.1) |
| *Wy. aporonoma*<br>[35] |  |  | **0.005**<br>(8.1) | **<0.0001**<br>(34.7) | 0.08 (3.0) |  |  |  |  |
| *Ae. albopictus*<br>[28] | **0.0001**<br>(20.6) | **0.0008**<br>(11.2) |  |  |  |  |  |  |  |
| *Sa. cyaneus*<br>[38] | **0.0001**<br>(20.8) |  |  |  |  |  | 0.05 (3.8) |  |  |
| *Li. pseudomethysticus* [14] | 0.02<br>(10.2) | **<0.0001**<br>(15.8) | 0.01 (6.7) | 0.02 (5.5) |  | 0.04 (4.1) | **0.005**<br>(7.8) | **0.004**<br>(8.4) |  |

\*Number of person days during which a species occurred (71 rainy and 75 dry season days, 2 x people sampling).
†Red and blue shaded cells indicate that a species was more common at ground level or on 5 m platforms, respectively.
‡Red and blue shaded cells indicate a negative (increase in occurrence) or positive (decrease in occurrence) association with mean weather, respectively.

Height and other environmental variables were included in both nominal logistic regression and standard least squares (Supplementary Table 6) models to assess their impact on species occurrence and abundance, respectively. During the rainy season, these models consistently indicated that *Hg. janthinomys* and *Sa. chloropterus* were more common on 5 m platforms than at ground level, while the remaining species, except for *Ps. amazonica* and *Sa. cyaneus*, which exhibited no height preference, were more common at ground level. Focusing primarily on highly significant variables or on all significant variables where occurrence and abundance models concurred, *Hg. janthinomys* was positively associated with mean temperature, while *Ps. amazonica* exhibited a negative correlation with the same variable. Both *Sabethes* species were negatively associated with mean weather, suggesting they are more active under clearer skies. An increase in 7-day cumulative rainfall lagged at 1 week was linked to a marginal decrease in *Hg. janthinomys* occurrence, as well as *Sa. cyaneus* occurrence and abundance, and an increase in *Li. durhamii* occurrence. Increasing rainfall lagged at 3 or 4 weeks was associated with an increase in occurrence or abundance of *Ps. amazonica*.

Supplementary analysis using a generalized linear model revealed interaction effects between edge type and height for occurrence of *Hg. janthinomys* and *Sa. chloropterus* sampled during the dry season (Supplementary Table 7). Simple effects tests for *Hg. janthinomys* showed a significant effect of edge type at ground level (Kruskal-Wallis Rank Sum, DF = 3, χ^2^ = 10.2, P = 0.02), where occurrence was highest at treefall gaps, and at 5 m (DF = 3, χ^2^ = 9.58, P = 0.02), where occurrence was highest at treefall gaps and in continuous forest. For *Sa. chloropterus*, there was no significant effect of edge type at ground level (DF = 3, χ^2^ = 2.51, P = 0.47), but a marginal effect at 5 m (DF = 3, χ^2^ = 7.35, P = 0.06), where occurrence was lower at urban edges than other edge types. The influence of these variables on species occurrence otherwise remained fairly consistent between seasons, although *Ps. amazonica* was marginally more common at ground level during the dry season. Furthermore, relative humidity had a greater impact on occurrence in the dry season, particularly for *Sa. chloropterus* and *Wy. aporonoma*, which exhibited positive associations with this variable. Mean weather was positively correlated with *Ps. amazonica* occurrence during the dry season but not during the rainy season, indicating its increased presence during harsher conditions. The same variable retained a negative association with *Sa. chloropterus*, confirming its heightened activity under clearer skies. Cumulative rainfall showed stronger positive associations with *Ps. amazonica*, reliant on ground water for breeding, and negative associations with *Li. pseudomethysticus*, during the dry season.

### Landscape context influenced changes in the vertical stratification of Sabethes subgenera

The two main Sabethes subgenera, *Sabethes* (N = 354) and *Sabethoides* (N = 257), differed in their vertical stratification. Contingency table analyses revealed a marginally lower overall occurrence of the subgenus *Sabethes* at 0 m compared to 5 m (Pearson’s chi-square, DF = 1, χ^2^ = 3.72, P = 0.054), and a substantially lower occurrence of *Sabethoides* at 0 m compared to 5 m (DF = 1, χ^2^ = 131, P < 0.0001) (Supplementary Table 8). When analyzed by edge type, the occurrence of both subgenera was lower at 0 m relative to 5 m within continuous forest (P < 0.0001 for both comparisons). At treefall gaps and rural edges, there was no significant difference in *Sabethes* occurrence between heights (P > 0.05 for both comparisons), although *Sabethoides* occurrence remained significantly lower at 0 m compared to 5 m (P < 0.0001 for both comparisons). Despite this, the ratio of *Sabethoides* occurring at 0 m and 5 m was 1:3.4 at rural edges compared to 1:16.4 in continuous forest. At urban edges, there was no notable difference in occurrence of either subgenus between heights (P > 0.2 for both comparisons), although *Sabethoides* mosquitoes were uncommon.

## Discussion

Landscape context may determine the role of forest edges in facilitating or retarding spillover and spillback of arboviruses. Our study shows that mosquito communities at edges bordering rural land cover are especially diverse and provide suitable refuge for known urban and sylvatic vectors. In contrast, those bordering urban land cover exhibit a reduced diversity and are less suitable for sylvatic species. However, urban edges intersect with the distribution of *Ae. albopictus* providing a pathway for its spread into forests^18^. Crucially, both anthropogenic and natural forest edges impact the vertical stratification of certain canopy-dwelling species, bringing them into contact with novel hosts.

In our study, mosquito communities were similar at treefall gaps and in continuous forest, while composition at rural edges was intermediate between interior forest and urban edges. Our findings are consistent with studies of plant communities showing that increased habitat contrast at forest edges negatively impacts suitability for specialist forest species^27,50^. In agreement, we sampled more forest mosquitoes at rural edges than at urban edges, which were dominated by ground dwelling *Limatus* and *Aedes* species. A loss of large trees^28^, positively associated with urbanization^51^, is likely to contribute to a decline in forest specialists, particularly those relying upon tree holes for breeding^52^. Our findings also revealed that ground level mosquito communities at treefall gaps were similar to elevated interior forest communities, whereas ground level continuous forest communities were more distinct. This suggests that canopy disruption impacts the vertical distribution of certain canopy-dwelling mosquito species, potentially bringing them into contact with terrestrial mammals associated with treefall gaps, including agoutis^53^, which have shown evidence of exposure to YFV^54^. Additionally, forest disturbance affects the vertical stratification of other canopy-dwelling wildlife^55,56^, although further studies are needed to determine the impact of edges on the distribution of vertebrates potentially involved in arbovirus transmission.

Species diversity was also shaped by landscape context, with mean species evenness highest at rural edges followed by continuous forest and lowest at treefall gaps and urban edges. This pattern persisted at ground level, but evenness decreased at higher elevations in continuous forest due to the dominance of *Hg. janthinomys*. These findings support the hypothesis of higher biodiversity and mixing of species from adjacent habitats at forest edges^57,58^, with the caveat that diversity of anthropophilic mosquitoes is higher at rural edges compared to urban edges. Rural edges also appear to enhance permeability for sylvatic vectors^8^ and inevitably their pathogens. Studies failing to detect higher diversity at forest edges have often restricted sampling to within a few hundred meters of the boundary^9–11,59^, yet we have only detected substantially lower diversity when sampling beyond 500 m into the forest^17,18^. We also found that species evenness was slightly, but significantly lower during the rainy season compared to the dry season. While higher diversity is generally associated with rainy months^11^, our findings may reflect an increased dominance of *Hg. janthinomys*, *Ps. amazonica*, and *Li. durhamii* during this period. Rarefaction estimates of species richness revealed less pronounced differences between forest edges, although these were based on the richness per edge type, which masked the heterogeneity of the urban edge sites. The heterogeneity of sampling sites at anthropogenic forest edges, influenced by neighboring habitat, can affect arthropod composition^60^.

It is well-established that forest edges significantly alter microclimate^17^, and our findings reflect this, with anthropogenic and natural edges exhibiting hotter and drier conditions compared to continuous forest. Landscape context also affected fluctuations in microclimate across the daily sampling hours, most notably at treefall gaps where temperature peaked, and relative humidity reached its lowest point in the early afternoon hours, as documented in our previous work^33^. We saw greater variation in microclimate between sites at urban edges (clustered in the southwest of the reserve) during the rainy season compared to the dry season, while the opposite was true at rural edges, treefall gaps, and in continuous forest (clustered in the northwest). Geographic orientation of sites, and season, are among the factors known to affect microclimate at the edges of forest fragments, with north-facing edges exhibiting higher temperatures and lower humidity than south-facing edges in the southern hemisphere^61^. These differences in microclimate may influence mosquito development rate^62^, pathogen extrinsic incubation period^63^, and other factors affecting vector competence^64^.

The means by which sylvatic mosquito-borne viruses are maintained throughout dry seasons has long intrigued researchers. Studies conducted in Panama during the 1950s, in tropical deciduous forest/ rainforest, showed reductions in *Haemagogus* and *Sabethes* abundance to very low levels during the dry season^65^. Our analysis revealed modest effects of season on mosquito abundance, with a tendency for higher numbers during the rainy season. However, our finding that key species persisted at appreciable levels during dry season months has important epidemiological implications. While transovarial transmission is often proposed as a mechanism for sustaining virus circulation^66^, we demonstrate that adult mosquito populations may play a crucial role in the Amazon, particularly in the forest canopy.

Environmental factors associated with the occurrence and abundance of key vector species can be used to characterize suitable habitat and refine risk models for pathogen emergence. We previously detected positive associations between *Haemagogus* mosquitoes, and both temperature and 7-day cumulative rainfall lagged at 1 week^32,33^. On this occasion, we confirmed the positive association between *Hg. janthinomys* and mean temperature but detected a marginal negative association with rainfall lagged at 1 week during the rainy season. Relationships between precipitation and tree hole breeding mosquitoes are likely to be complex. While rainfall is essential for eggs to hatch, too much water can flush out mosquitoes and diminish populations^67^. This species was seldom encountered at anthropogenic edges. It was mostly active inside the forest and above the ground but descended to ground level in abundance at treefall gaps, making *Hg. janthinomys* a potential bridge vector in these settings. Research is now required to assess the vector competence of this species for DENV, ZIKV, CHIKV, and MAYV^68,69^.

We have sampled *Ps. amazonica* in high relative abundance throughout our studies^17,18,32,33^. It was mainly captured in continuous forest and at treefall gaps but was also the most abundant mosquito at rural edges. Again, its relationship with precipitation is likely to be complex, as indicated by associations with rainfall that deviated from our previous results^32^. Species of the subgenus *Janthinosoma* lay desiccation resistant eggs in pools of ground water^45,70^. At the Ducke reserve, these only form after sustained heavy rainfall saturates the forest floor. However, once established, *Psorophora* species can develop through multiple generations very quickly^71^. We detected a negative association with temperature in the rainy season adding to evidence that *Ps. amazonica* tolerates cool conditions^33^. There was a limited effect of height on the occurrence and abundance of *Ps. amazonica*, which has potential to interact with humans and with wildlife at ground level and in the canopy. Despite these traits and its aggressive biting behavior, we know nothing about its vector status. However, other closely related *Janthinosoma* species harbor medically important orthoflaviviruses including Ilhéus and West Nile viruses^70^.

*Sabethes* mosquitoes, recognized as important secondary vectors of YFV^72^, were common in both continuous forest and at treefall gaps, particularly under favorable weather coniditions. Notably, *Sa. chloropterus* was frequently captured at rural edges, further emphasizing the affinity of certain *Sabethes* species for edge habitats^17,52,73^. However, the contrasting vertical distributions of *Sa.* (*Sabethoides*) *chloropterus* and *Sa.* (*Sabethes*) *cyaneus* has important implications for pathogen transmission. In line with Galindo et al.^74^, our findings revealed a strong preference for elevated heights by *Sa. chloropterus*, while *Sa. cyaneus* only exhibited a slight, insignificant, preference. Our subgenus analysis confirmed these observations and demonstrated a relative increase in *Sabethoides* mosquitoes at ground level at rural edges compared to continuous forest. *Sabethes* species possess a strong bridge vector potential and should be targeted for arbovirus surveillance, including with BG-Sentinel traps^17,32^. These mosquitoes display an intriguing preference for biting noses^75^, a behavior that could be exploited in the development of attractants.

The Asian tiger mosquito, *Ae. albopictus*, has gained attention for its potential role as a bridge vector^76^, which is unsurprising considering its presence at forest edges, global distribution, and critical vector status^5^. After *Li. durhamii*, *Ae. albopictus* was the most abundant species found at urban edges. Neither species was common at 5 m indicating that their role as a bridge vector might be confined close to ground level. Other studies investigating *Ae. albopictus* activity have also shown that this species is mainly active near the ground^77,78^. However, our understanding of the vertical stratification of *Ae. albopictus* at forest edges remains limited, and further research that considers the broader landscape context is needed. Whereas *Ae. albopictus* is widely recognized as an important global vector^1,5^, evidence of a role for *Li. durhamii* in arbovirus transmission is scarce^79^.

While our study provides valuable insights into mosquito dynamics at forest edges, there were several limitations. The small number of sampling sites at each edge type may restrict the generalizability of findings, yet the tantalizing heterogeneity of mosquito communities among urban edge sites deserves further investigation. Furthermore, our sampling strategy focused on peak activity times of major sylvatic vectors^33^ which may have underestimated the abundance of species like *Ae. albopictus*, if exhibiting different activity patterns^43^. Simultaneous ground and platform sampling could underestimate vertical mosquito movement, but rotating sampling introduces logistical challenges. Lastly, a single-year study may not capture the full range of mosquito community dynamics. Sustained, multi-year investigations to assess the influence of annual environmental variations are needed.

The landscape context of forest edges must be considered when assessing pathogen emergence risk, along with relative human population densities, and interactions between humans and forest environments. A synergistic approach integrating in-depth field studies with big data analysis will be crucial to understanding the nuances of human-mosquito-wildlife interactions and developing risk models that accurately reflect the dynamics of these complex ecological systems.

## Acknowledgements

We wish to acknowledge Raimundo Coelho and Aleksey Lima (FMT-HVD) for driving and logistics, and the Instituto Nacional de Pesquisas da Amazônia (INPA) for permission to use the Adolpho Ducke Reserve to collect samples for this study.

## Author contributions

Conceptualization, A.H. and K.A.H.; Data curation, A.H., I.P., A.G., T.N.F.S., and C.R.M.; Formal analysis, A.H, V.S., and K.A.H.; Funding acquisition, M.V.G.d.L., N.V., and K.A.H.; Investigation, A.H., N.F.F., I.P., A.G., T.N.F.S., C.R.M., J.T.A.J., F.P.A., and E.R.C.; Methodology, A.H., J.T.A.J., F.P.A., E.R.C., M.P.G.M., and K.A.H.; Supervision, M.G., M.V.G.d.L., M.P.G.M., N.V., and K.A.H.; Visualization, A.H., V.S., M.B., and K.A.H.; Writing—original draft, A.H. and K.A.H.; Writing—review and editing, A.H., V.S., M.G., V.M.S., M.B., M.V.G.d.L., M.P.G.M., N.V., and K.A.H. All authors have read and agreed to the published version of the manuscript.

## Data availability

All data generated or analyzed during this study are available in the Mendeley Data repository, DOI: 10.17632/nyjjmc2htd.1

## References

1 Weaver, S. C., Charlier, C., Vasilakis, N. & Lecuit, M. Zika, Chikungunya, and Other Emerging Vector-Borne Viral Diseases. Annu. Rev. Med. 69, 395–408, doi:10.1146/annurev-med-050715-105122 (2018).

2 Guth, S., Hanley, K. A., Althouse, B. M. & Boots, M. Ecological processes underlying the emergence of novel enzootic cycles: Arboviruses in the neotropics as a case study. PLoS Negl. Trop. Dis. 14, e0008338, doi:10.1371/journal.pntd.0008338 (2020).

3 LaBeaud, A., Bashir, F. & King, C. H. Measuring the burden of arboviral diseases: the spectrum of morbidity and mortality from four prevalent infections. Popul. Health Metr. 9, 1, doi:10.1186/1478-7954-9-1 (2011).

4 Puntasecca, C. J., King, C. H. & LaBeaud, A. D. Measuring the global burden of chikungunya and Zika viruses: A systematic review. PLoS Negl. Trop. Dis. 15, e0009055, doi:10.1371/journal.pntd.0009055 (2021).

5 Wilder-Smith, A. et al. Epidemic arboviral diseases: priorities for research and public health. Lancet Infect. Dis. 17, e101–e106, doi:10.1016/S1473-3099(16)30518-7 (2017).

6 Barrett, A. D. & Higgs, S. Yellow fever: a disease that has yet to be conquered. Annu. Rev. Entomol. 52, 209–229, doi:10.1146/annurev.ento.52.110405.091454 (2007).

7 Bryant, J. E., Holmes, E. C. & Barrett, A. D. Out of Africa: a molecular perspective on the introduction of yellow fever virus into the Americas. PLoS Pathog. 3, e75, doi:10.1371/journal.ppat.0030075 (2007).

8 Borremans, B., Faust, C., Manlove, K. R., Sokolow, S. H. & Lloyd-Smith, J. O. Cross-species pathogen spillover across ecosystem boundaries: mechanisms and theory. Philos. Trans. R. Soc. Lond. B Biol. Sci. 374, 20180344, doi:10.1098/rstb.2018.0344 (2019).

9 Steiger, D. M., Ritchie, S. A. & Laurance, S. G. W. Mosquito communities and disease risk influenced by land use change and seasonality in the Australian tropics. Parasit. Vectors. 9, 387, doi:10.1186/s13071-016-1675-2 (2016).

10 Young, K. I., Buenemann, M., Vasilakis, N., Perera, D. & Hanley, K. A. Shifts in mosquito diversity and abundance along a gradient from oil palm plantations to conterminous forests in Borneo. Ecosphere 12, e03463, doi:10.1002/ecs2.3463 (2021).

11 Costa, L. N. P., Novais, S., Oki, Y., Fernandes, G. W. & Borges, M. A. Z. Mosquito (Diptera: Culicidae) diversity along a rainy season and edge effects in a riparian forest in Southeastern Brazil. Austral Ecol. 48, 41–55, doi:10.1111/aec.13250 (2023).

12 Almeida, J. F. et al. Change in the faunal composition of mosquitoes (Diptera: Culicidae) along a heterogeneous landscape gradient in the Brazilian Amazon. PLoS One 18, e0288646, doi:10.1371/journal.pone.0288646 (2023).

13 Suzán, G. et al. Epidemiological Considerations of Rodent Community Composition in Fragmented Landscapes in Panama. J. Mammal. 89, 684–690, doi:10.1644/07-MAMM-A-015R1.1 (2008).

14 Villaseñor, N. R., Driscoll, D. A., Escobar, M. A. H., Gibbons, P. & Lindenmayer, D. B. Urbanization Impacts on Mammals across Urban-Forest Edges and a Predictive Model of Edge Effects. PLoS One 9, e97036, doi:10.1371/journal.pone.0097036 (2014).

15 Mendes-Oliveira, A. C. et al. Oil palm monoculture induces drastic erosion of an Amazonian forest mammal fauna. PLoS One 12, e0187650, doi:10.1371/journal.pone.0187650 (2017).

16 Steiger, D. M. et al. Effects of landscape disturbance on mosquito community composition in tropical Australia. J. Vector Ecol. 37, 69–76, doi:10.1111/j.1948-7134.2012.00201.x (2012).

17 Hendy, A. et al. Where boundaries become bridges: Mosquito community composition, key vectors, and environmental associations at forest edges in the central Brazilian Amazon. PLoS Negl. Trop. Dis. 17, e0011296, doi:10.1371/journal.pntd.0011296 (2023).

18 Hendy, A. et al. Into the woods: Changes in mosquito community composition and presence of key vectors at increasing distances from the urban edge in urban forest parks in Manaus, Brazil. Acta Trop. 206, 105441, doi:10.1016/j.actatropica.2020.105441 (2020).

19 Lenz, B. B., Jack, K. M. & Spironello, W. R. Edge effects in the primate community of the biological dynamics of Forest Fragments Project, Amazonas, Brazil. Am. J. Phys. Anthropol. 155, 436–446, doi:10.1002/ajpa.22590 (2014).

20 Mittermeier, R. A. & van Roosmalen, M. G. Preliminary observations on habitat utilization and diet in eight Surinam monkeys. Folia Primatol. (Basel) 36, 1–39, doi:10.1159/000156007 (1981).

21 Bolt, L. M. et al. The influence of anthropogenic edge effects on primate populations and their habitat in a fragmented rainforest in Costa Rica. Primates 59, 301–311, doi:10.1007/s10329-018-0652-0 (2018).

22 Sobroza, T. V. et al. Does co-occurrence drive vertical niche partitioning in parapatric tamarins (*Saguinus* spp.)? Austral Ecol. 46, 1335–1342, doi:10.1111/aec.13085 (2021).

23 Gordo, M., Calleia, F. O., Vasconcelos, S. A., Leite, J. J. F. & Ferrari, S. F. in Primates in Fragments (eds L. Marsh & C. Chapman) 357-370 (Springer, 2013).

24 Boyle, S. A. et al. in Amazonian Mammals: Current Knowledge and Conservation Priorities (eds Wilson R. Spironello et al.) Ch. 13, 335-363 (Springer International Publishing, 2023).

25 dos Santos, L. S., Pereira, H. d. S. & Gordo, M. Simpatria entre populações humanas e de sauim- de-coleira (*Saguinus bicolor*) em fragmentos florestais de Manaus, Amazonas. Neotrop. Primates 23, 25–32, doi:10.62015/np.2017.v23.119 (2023).

26 Câmara, D. C. P., Pinel, C. d. S., Rocha, G. P., Codeço, C. T. & Honório, N. A. Diversity of mosquito (Diptera: Culicidae) vectors in a heterogeneous landscape endemic for arboviruses. Acta Trop. 212, 105715, doi:10.1016/j.actatropica.2020.105715 (2020).

27 Guerra, T. N. F., Araújo, E. L., Sampaio, E. V. S. B. & Ferraz, E. M. N. Urban or rural areas: which types of surrounding land use induce stronger edge effects on the functional traits of tropical forests plants? Appl. Veg. Sci. 20, 538–548, doi:10.1111/avsc.12315 (2017).

28 Laurance, W. F., Delamônica, P., Laurance, S. G., Vasconcelos, H. L. & Lovejoy, T. E. Rainforest fragmentation kills big trees. Nature 404, 836–836, doi:10.1038/35009032 (2000).

29 Roosmalen, M. G. M. v. Habitat preferences, diet, feeding strategy and social organization of the black spider monkey [*Ateles paniscus paniscus* Linnaeus 1758] in Surinam. Acta Amaz. 15, 7–238, doi:10.1590/1809-43921985155238 (1985).

30 Galindo, P., Carpenter, S. J. & Trapido, H. Ecological observations on forest mosquitoes of an endemic yellow fever area in Panama. Am. J. Trop. Med. Hyg. 31, 98–137, doi:10.4269/ajtmh.1951.s1-31.98 (1951).

31 Magura, T., Lövei, G. L. & Tóthmérész, B. Edge responses are different in edges under natural versus anthropogenic influence: a meta-analysis using ground beetles. Ecol. Evol. 7, 1009–1017, doi:doi.org/10.1002/ece3.2722 (2017).

32 Hendy, A. et al. The vertical stratification of potential bridge vectors of mosquito-borne viruses in a central Amazonian forest bordering Manaus, Brazil. Sci. Rep. 10, 18254, doi:10.1038/s41598-020-75178-3 (2020).

33 Hendy, A. et al. Microclimate and the vertical stratification of potential bridge vectors of mosquito-borne viruses captured by nets and ovitraps in a central Amazonian forest bordering Manaus, Brazil. Sci. Rep. 11, 21129, doi:10.1038/s41598-021-00514-0 (2021).

34 Oliveira, M. L. d., Baccaro, F. B., Braga-Neto, R. & Magnusson, W. E. Reserva Ducke: A bioversidade Amazônica através de uma grade. (Instituto Nacional de Pesquisas da Amazônia, 2011).

35. Gordo, M., Rodrigues, L. F. R., Vidal, M. D., Spironello, W. R. & Ribeiro, F. R. P. in Reserva Ducke: A bioversidade Amazônica através de uma grade (eds M. L. de Oliveira, F. B. Baccaro, R. Braga- Neto, & W. E. Magnusson) 39-49 (Instituto Nacional de Pesquisas da Amazônia, 2011).

36 Carvalho, T. P. V. d. & Costa, R. C. in 2013: *IV Congreso Latinoamericano de Agroecología*. (Sociedad Científica Latinoamericana de Agroecología).

37 Gonçalves Maciel, L. H., et al. Prevalence of arboviruses and other infectious causes of skin rash in patients treated at a tertiary health unit in the Brazilian Amazon. PLoS Negl. Trop. Dis. 16, e0010727, doi:10.1371/journal.pntd.0010727 (2022).

38 Saraiva, M. d. G. G., et al. Historical analysis of the records of sylvan yellow fever in the State of Amazonas, Brazil, from 1996 to 2009. *Rev. Soc. Bras. Med. Trop.* **46**, 223-226, doi:10.1590/0037-8682-1573-2013 (2013).

39 Mourão, M. P. et al. Arboviral diseases in the Western Brazilian Amazon: A perspective and analysis from a tertiary health & research center in Manaus, State of Amazonas. Rev. Soc. Bras. Med. Trop. 48, 20–26, doi:10.1590/0037-8682-0133-2013 (2015).

40. United States Geological Survey. Earth Explorer, <https://earthexplorer.usgs.gov/> (2017).

41. GISMAPS. Districts of Manaus (SHP), <https://gismaps.com.br/en/downloads/bairros-de-manaus-shp/> (2023).

42 Trapido, H., Galindo, P. & Carpenter, S. J. A survey of forest mosquitoes in relation to sylvan yellow fever in the Panama Isthmian area. Am. J. Trop. Med. Hyg. 4, 525–542, doi:10.4269/ajtmh.1955.4.525 (1955).

43 Hawley, W. A. The biology of *Aedes albopictus*. J. Am. Mosq. Control Assoc. Suppl. 1, 1–39 (1988).

44 Instituto Nacional de Meteorologia. Banco de dados meteorológicos para ensino e pesquisa, <https://portal.inmet.gov.br/> (2023).

45. Wilkerson, R. C., Linton, Y.-M. & Strickman, D. Mosquitoes of the World. Vol. 1 and 2 (Johns Hopkins University Press, 2021).

46 JMP 17.1.0 (SAS Institute Inc, Cary, NC, USA, 2023).

47 Hammer, Ø., Harper, D. A. T. & Ryan, P. D. PAST: Paleontological statistics software package for education and data analysis. *Palaeontol*. Electronica 4, 9 (2001).

48 Hsieh, T. C., Ma, K. H. & Chao, A. iNEXT: an R package for rarefaction and extrapolation of species diversity (Hill numbers). Methods Ecol. Evol. 7, 1451–1456, doi:10.1111/2041-210X.12613 (2016).

49 Hendy, A., et al. Dataset: Forest edge landscape context affects mosquito community composition and risk of pathogen emergence. Mendeley Data, doi:10.17632/nyjjmc2htd.1 (2024).

50 Vallet, J., Beaujouan, V., Pithon, J., Rozé, F. & Daniel, H. The effects of urban or rural landscape context and distance from the edge on native woodland plant communities. Biodivers. Conserv. 19, 3375–3392, doi:10.1007/s10531-010-9901-2 (2010).

51 Wang, Z. & Yang, J. Urbanization strengthens the edge effects on species diversity and composition of woody plants in remnant forests. For. Ecosyst. 9, 100063, doi:10.1016/j.fecs.2022.100063 (2022).

52 Mangudo, C., Aparicio, J. P., Rossi, G. C. & Gleiser, R. M. Tree hole mosquito species composition and relative abundances differ between urban and adjacent forest habitats in northwestern Argentina. Bull. Entomol. Res. 108, 203–212, doi:10.1017/s0007485317000700 (2018).

53 Dubost, G. Ecology and social life of the red acouchy, *Myoprocta exilis*; comparison with the orange-rumped agouti, *Dasyprocta leporina*. J. Zool. 214, 107–123, doi:10.1111/j.1469-7998.1988.tb04990.x (1988).

54 de Thoisy, B., Dussart, P. & Kazanji, M. Wild terrestrial rainforest mammals as potential reservoirs for flaviviruses (yellow fever, dengue 2 and St Louis encephalitis viruses) in French Guiana. Trans. R. Soc. Trop. Med. Hyg. 98, 409–412, doi:10.1016/j.trstmh.2003.12.003 (2004).

55 Tregidgo, D. J., Qie, L., Barlow, J., Sodhi, N. S. & Lim, S. L.-H. Vertical Stratification Responses of an Arboreal Dung Beetle Species to Tropical Forest Fragmentation in Malaysia. Biotropica 42, 521–525, doi:10.1111/j.1744-7429.2010.00649.x (2010).

56 Silva, I., Rocha, R., López-Baucells, A., Farneda, F. Z. & Meyer, C. F. J. Effects of Forest Fragmentation on the Vertical Stratification of Neotropical Bats. Diversity 12, 67 (2020).

57 Duelli, P., Obrist, M. K. & Flückiger, P. F. Forest edges are biodiversity hotspots - also for Neuroptera. Acta Zool. Acad. Sci. Hung. 48, 75–87 (2002).

58 Baker, J., French, K. & Whelan, R. J. The edge effect and ecotonal species: Bird communities across a natural edge in southeastern Australia. Ecology 83, 3048–3059, doi:10.1890/0012-9658(2002)083[3048:TEEAES]2.0.CO;2 (2002).

59 Reiskind, M. H., Griffin, R. H., Janairo, M. S. & Hopperstad, K. A. Mosquitoes of field and forest: the scale of habitat segregation in a diverse mosquito assemblage. Med. Vet. Entomol. 31, 44–54, doi:10.1111/mve.12193 (2017).

60 Magura, T. Carabids and forest edge: spatial pattern and edge effect. For. Ecol. Manage. 157, 23–37, doi:10.1016/S0378-1127(00)00654-X (2002).

61 Bernaschini, M. L., Trumper, E., Valladares, G. & Salvo, A. Are all edges equal? Microclimatic conditions, geographical orientation and biological implications in a fragmented forest. Agric. Ecosyst. Environ. 280, 142–151, doi:10.1016/j.agee.2019.04.035 (2019).

62 Bates, M. The development and longevity of *Haemagogus* mosquitoes under laboratory conditions. Ann. Entomol. Soc. Am. 40, 1–12, doi:10.1093/aesa/40.1.1 (1947).

63 Bates, M. & Roca-Garcia, M. The development of the virus of yellow fever in *Haemagogus* mosquitoes. Am. J. Trop. Med. Hyg. 26, 585–605, doi:10.4269/ajtmh.1946.s1-26.585 (1946).

64 Mordecai, E. A. et al. Thermal biology of mosquito-borne disease. Ecol. Lett. 22, 1690–1708, doi:10.1111/ele.13335 (2019).

65 Galindo, P., Trapido, H., Carpenter, S. J. & Blanton, F. S. The abundance cycles of arboreal mosquitoes during six years at a sylvan yellow fever locality in Panama. Ann. Entomol. Soc. Am. 49, 543–547, doi:10.1093/aesa/49.6.543 (1956).

66 Dutary, B. E. & Leduc, J. W. Transovarial transmission of yellow fever virus by a sylvatic vector, *Haemagogus equinus*. Trans. R. Soc. Trop. Med. Hyg. 75, 128 (1981).

67 Washburn, J. O. & Anderson, J. R. Habitat overflow, a source of larval mortality for *Aedes sierrensis* (Diptera: Culicidae). J. Med. Entomol. 30, 802–804, doi:10.1093/jmedent/30.4.802 (1993).

68 Hendy, A. et al. Towards the Laboratory Maintenance of Haemagogus janthinomys (Dyar, 1921), the Major Neotropical Vector of Sylvatic Yellow Fever. *Viruses* **15**, 45 (2023).

69 Azar, S. R. & Weaver, S. C. Vector Competence Analyses on *Aedes aegypti* Mosquitoes using Zika Virus. JoVE, e61112, doi:10.3791/61112 (2020).

70 Mello, C. F., Santos-Mallet, J. R., Tátila-Ferreira, A. & Alencar, J. Comparing the egg ultrastructure of three *Psorophora ferox* (Diptera: Culicidae) populations. Braz. J. Biol. 78, 505–508, doi:10.1590/1519-6984.171829 (2018).

71 Harrison, B. A., Varnado, W., Whitt, P. B. & Goddard, J. New diagnostic characters for females of *Psorophora* (*Janthinosoma*) species in the United States, with notes on *Psorophora mexicana* (Bellardi) (Diptera: Culicidae). J. Vector Ecol. 33, 232–237, doi:10.3376/1081-1710-33.2.232 (2008).

72 Abreu, F. V. S., et al. Haemagogus leucocelaenus and Haemagogus janthinomys are the primary vectors in the major yellow fever outbreak in Brazil, 2016-2018. *Emerg. Microbes. Infect.* **8**, 218- 231, doi:10.1080/22221751.2019.1568180 (2019).

73 Almeida, J. F., Belchior, H. C. M., Ríos-Velásquez, C. M. & Pessoa, F. A. C. Diversity of mosquitoes (Diptera: Culicidae) collected in different types of larvitraps in an Amazon rural settlement. PLoS One 15, e0235726, doi:10.1371/journal.pone.0235726 (2020).

74 Galindo, P., Trapido, H. & Carpenter, S. J. Observations on diurnal forest mosquitoes in relation to sylvan yellow fever in Panama. Am. J. Trop. Med. Hyg. 30, 533–574, doi:10.4269/ajtmh.1950.s1-30.533 (1950).

75 Dégallier, N., Sá Filho, G. C., Silva, O. V. & Travassos da Rosa, A. P. A. Comportamento de pouso sobre partes do corpo humano, em mosquitos da floresta amazonica (Diptera : Culicidae). *Bol. Mus. Para. Emílio Goeldi*, Zoo. 6, 97–108 (1990).

76 Pereira-dos-Santos, T., Roiz, D., Lourenço-de-Oliveira, R. & Paupy, C. A Systematic Review: Is *Aedes albopictus* an Efficient Bridge Vector for Zoonotic Arboviruses? Pathogens 9, 266, doi:10.3390/pathogens9040266 (2020).

77 Obenauer, P. J., Kaufman, P. E., Kline, D. L. & Allan, S. A. Detection of and monitoring for *Aedes albopictus* (Diptera: Culicidae) in suburban and sylvatic habitats in north central Florida using four sampling techniques. Environ. Entomol. 39, 1608–1616, doi:10.1603/en09322 (2010).

78 Dias, R., de Mello, C. F., Santos, G. S., Carbajal-de-la-Fuente, A. L. & Alencar, J. Vertical Distribution of Oviposition and Temporal Segregation of Arbovirus Vector Mosquitoes (Diptera: Culicidae) in a Fragment of the Atlantic Forest, State of Rio de Janeiro, Brazil. Trop. Med. Infect. Dis. 8, 256, doi:10.3390/tropicalmed8050256 (2023).

79 Barrio-Nuevo, K. M. et al. Detection of Zika and dengue viruses in wild-caught mosquitoes collected during field surveillance in an environmental protection area in São Paulo, Brazil. PLoS One 15, e0227239, doi:10.1371/journal.pone.0227239 (2020).

