## Supplementary Information for "Forest edge landscape context affects mosquito community composition and risk of pathogen emergence"

**Supplementary information**
Supplementary Figures 1 – 2, Supplementary Tables 1 – 8

### Forest edge landscape context affects mosquito community composition and risk of pathogen emergence

Adam Hendy[1,2], Nelson Ferreira Fé[3,†], Igor Pedrosa[3], André Girão[3], Taly Nayandra Figueira dos Santos[3], Claudia Reis Mendonça[3], José Tenaçol Andes Júnior[3], Flamarion Prado Assunção[3], Edson Rodrigues Costa[4], Vincent Sluydts[5], Marcelo Gordo[4], Vera Margarete Scarpassa[6], Michaela Buenemann[7], Marcus Vinícius Guimarães de Lacerda[1,3,8], Maria Paula Gomes Mourão[3], Nikos Vasilakis[1,9,10]* and Kathryn A. Hanley[2]*

[1]Department of Pathology, University of Texas Medical Branch, Galveston, Texas, USA
[2]Department of Biology, New Mexico State University, Las Cruces, New Mexico, USA
[3]Fundação de Medicina Tropical Doutor Heitor Vieira Dourado (FMT-HVD), Manaus, Amazonas, Brazil
[4]Laboratório de Biologia da Conservação, Projeto Sauim-de-Coleira, Instituto de Ciências Biológicas, Universidade Federal do Amazonas, Manaus, Amazonas, Brazil
[5]Department of Biology, University of Antwerp, Evolutionary Ecology Group, Wilrijk, Belgium
[6]Coordenação de Biodiversidade, Instituto Nacional de Pesquisas da Amazônia, Manaus, Amazonas, Brazil
[7]Department of Geography and Environmental Studies, New Mexico State University, Las Cruces, New Mexico, USA
[8]Instituto Leônidas & Maria Deane (Fiocruz - Amazônia), Manaus, Amazonas, Brazil
[9]Center for Vector-Borne and Zoonotic Diseases, University of Texas Medical Branch, Galveston, Texas, USA
[10]Institute for Human Infection and Immunity, University of Texas Medical Branch, Galveston, Texas, USA

[†]Deceased

**Supplementary Figure 1.** Sampling sites

**Supplementary Figure 1.** Sampling sites. A – C = continuous forest, D – E = treefall gap, G – I rural edge, and J – L = urban edge. The image at site E was taken in May 2023 during a revisit and shows some regrowth of vegetation near the forest floor. All other images were taken at the time of the study.

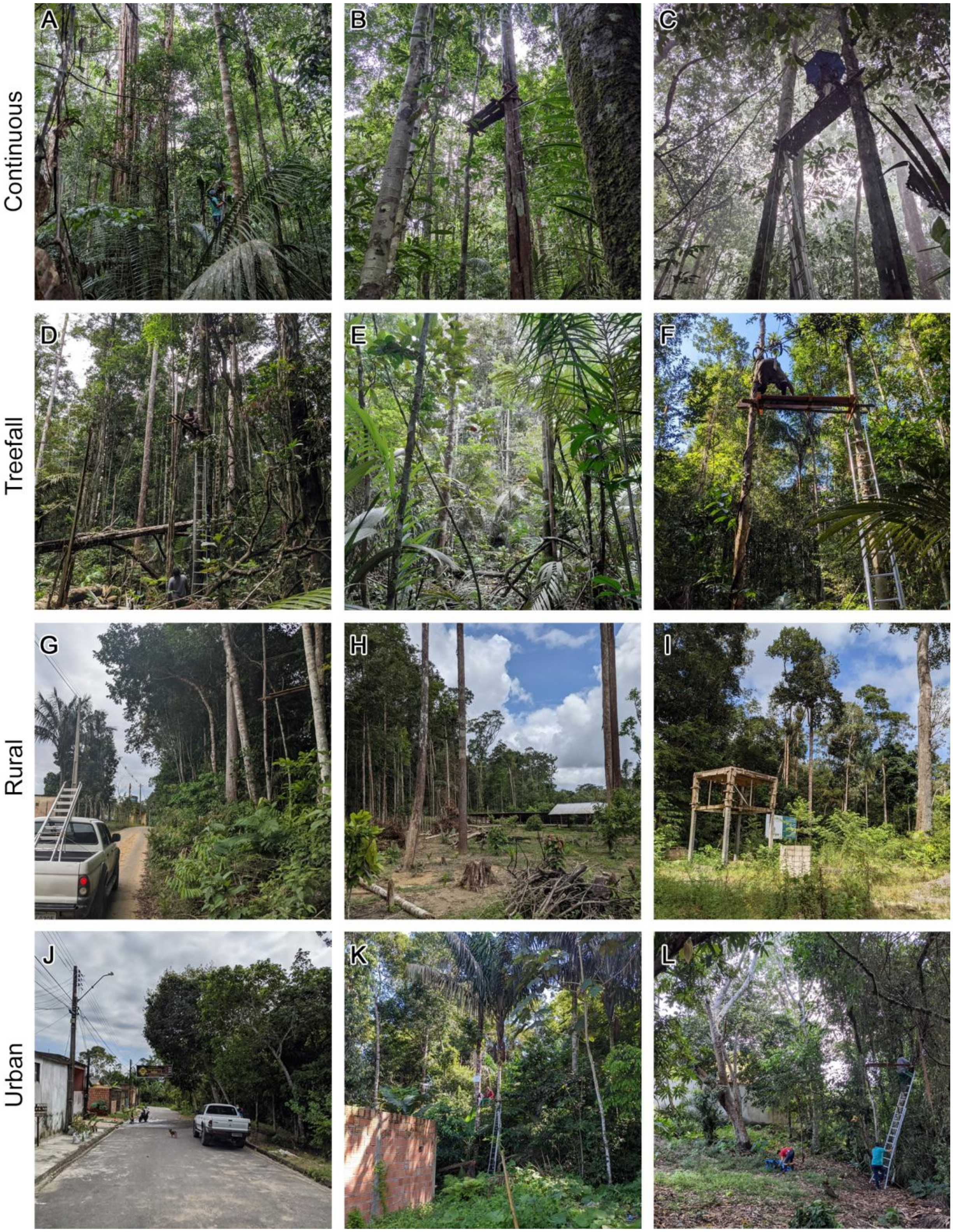

**Supplementary Figure 2.** Rarefaction analysis

**Supplementary Figure 2.** Species richness rarefaction and extrapolation curves. Upper panel shows species richness for data grouped by edge type. Lower panels show species richness for data grouped by edge type for each height (0 m and 5 m). Shaded areas surrounding rarefaction and extrapolation lines represent 95% confidence intervals.

*By edge type*

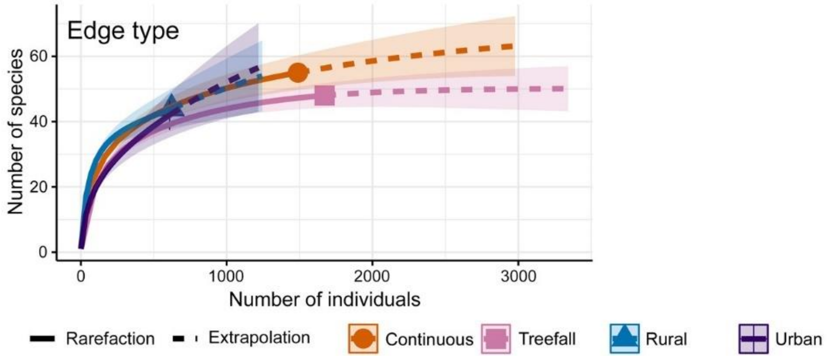

*By edge type and height*

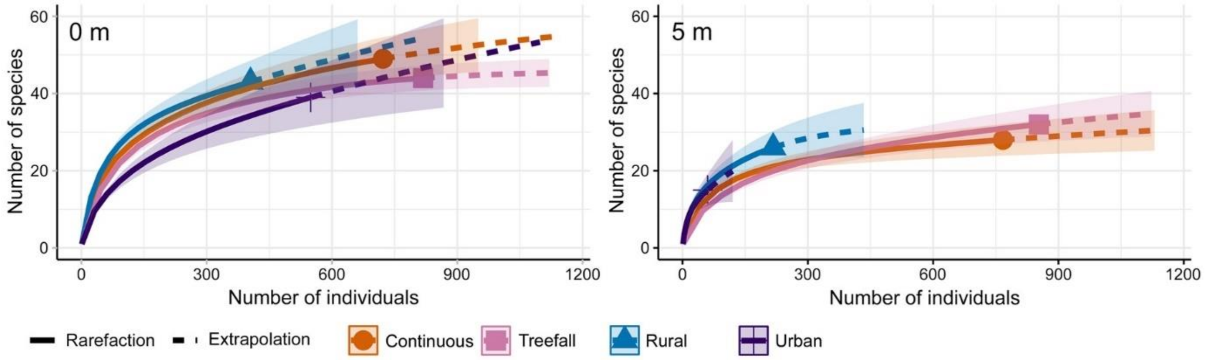

**Supplementary Table 1.** Morisita index

**Supplementary Table 1.** Morisita overlap index for various matrices of comparison. Cont. = Continuous forest.

*By site at each edge type*

| | **Cont. A** | **Cont. B** | **Cont. C** |
|---|---|---|---|
| **Cont. A** | 1 | | |
| **Cont. B** | 0.955 | 1 | |
| **Cont. C** | 0.928 | 0.967 | 1 |

| | **Treefall D** | **Treefall E** | **Treefall F** |
|---|---|---|---|
| **Treefall D** | 1 | | |
| **Treefall E** | 0.918 | 1 | |
| **Treefall F** | 0.975 | 0.962 | 1 |

| | **Rural G** | **Rural H** | **Rural I** |
|---|---|---|---|
| **Rural G** | 1 | | |
| **Rural H** | 0.561 | 1 | |
| **Rural I** | 0.823 | 0.720 | 1 |

| | **Urban J** | **Urban K** | **Urban L** |
|---|---|---|---|
| **Urban J** | 1 | | |
| **Urban K** | 0.285 | 1 | |
| **Urban L** | 0.893 | 0.589 | 1 |

*By edge type and season at 0 m* (R = rainy, D = dry)

| | **Cont. R** | **Treefall R** | **Rural R** | **Urban R** | **Cont. D** | **Treefall D** | **Rural D** | **Urban D** |
|---|---|---|---|---|---|---|---|---|
| **Cont. R** | 1 | | | | | | | |
| **Treefall R** | 0.852 | 1 | | | | | | |
| **Rural R** | 0.782 | 0.617 | 1 | | | | | |
| **Urban R** | 0.090 | 0.052 | 0.544 | 1 | | | | |
| **Cont. D** | 0.923 | 0.740 | 0.736 | 0.076 | 1 | | | |
| **Treefall D** | 0.779 | 0.951 | 0.540 | 0.061 | 0.703 | 1 | | |
| **Rural D** | 0.686 | 0.555 | 0.809 | 0.439 | 0.722 | 0.586 | 1 | |
| **Urban D** | 0.129 | 0.085 | 0.555 | 0.924 | 0.117 | 0.089 | 0.489 | 1 |

*By edge type and season at 5 m* (urban edge excluded due to small sample size)

| | **Cont. R** | **Treefall R** | **Rural R** | **Cont. D** | **Treefall D** | **Rural D** |
|---|---|---|---|---|---|---|
| **Cont. R** | 1 | | | | | |
| **Treefall R** | 0.985 | 1 | | | | |
| **Rural R** | 0.614 | 0.551 | 1 | | | |
| **Cont. D** | 0.975 | 0.950 | 0.652 | 1 | | |
| **Treefall D** | 0.991 | 0.974 | 0.606 | 0.986 | 1 | |
| **Rural D** | 0.376 | 0.313 | 0.769 | 0.491 | 0.433 | 1 |

**Supplementary Table 2.** PCA loadings

**Supplementary Table 2.** Loading matrices for principal components analysis. PC1 and PC2 were included in hierarchical clustering by edge type, and PC1, PC2, and PC3 were included in hierarchical clustering by edge type and height.

*By edge type*

| **Species** | **Prin1** | **Prin2** | **Prin3** |
|---|---|---|---|
| *Cx.* (*Mel.*) *caudelli* | 0.991311 | -0.120233 | 0.053350 |
| *Hg.* (*Hag.*) *janthinomys* | 0.950449 | -0.000153 | -0.310880 |
| *Sa.* (*Sab.*) *bipartipes* | -0.933055 | 0.319997 | 0.164349 |
| *Cx.* (*Mel.*) *adamesi* | 0.913308 | -0.098913 | -0.395076 |
| *Hg.* (*Con.*) *leucocelaenus* | 0.904867 | 0.298701 | 0.303305 |
| *Sa.* (*Sab.*) *belisarioi* | -0.827315 | 0.540744 | -0.152141 |
| *Sa.* (*Sab.*) *quasicyaneus* | 0.711878 | 0.382113 | -0.589253 |
| *Sa.* (*Sbo.*) *chloropterus* | -0.038068 | 0.995588 | 0.085761 |
| *Sa.* (*Sab.*) *batesi* | 0.216274 | 0.950570 | -0.222804 |
| *Wy. argenteorostris* | -0.286225 | 0.913786 | -0.288220 |
| *Ae.* (*Och.*) *hastatus* | -0.475948 | 0.878664 | 0.037722 |
| *Sa.* (*Sab.*) *albiprivus* | -0.349804 | 0.829002 | -0.436339 |
| *Wy.* (*Dod.*) *aphobema* | -0.122288 | 0.777124 | 0.617353 |
| *Sa.* (*Sab.*) *cyaneus* | 0.541808 | 0.774269 | 0.327035 |
| *Wy.* (*Wyo.*) *hemisagnosta* | -0.654374 | -0.755670 | 0.027520 |
| *Ae.* (*Stg.*) *aegypti* | -0.670203 | -0.742157 | -0.005546 |
| *Cx.* (*Cux.*) *coronator* | 0.411645 | -0.338475 | 0.846158 |
| *Wy.* (*Cru.*) *kummi* | 0.553051 | -0.044073 | -0.831981 |
| *Ae.* (*Och.*) *nubilus* | 0.593100 | -0.092434 | 0.799805 |
| *Cx.* (*Cux.*) *declarator* | 0.202945 | 0.579126 | 0.789574 |

Shaded cells have absolute loading values < ±0.3.

**Supplementary Table 2.** PCA loadings

*By edge type and height*

| Species | Prin1 | Prin2 | Prin3 | Prin4 | Prin5 | Prin6 | Prin7 |
|---|---|---|---|---|---|---|---|
| *Sa.* (*Sab.*) *cyaneus* | 0.874510 | 0.112199 | 0.012741 | 0.287447 | 0.356118 | 0.113833 | -0.008838 |
| *Wy.* (*Wyo.*) *hemisagnosta* | -0.766426 | -0.584430 | 0.156888 | 0.147692 | -0.091739 | 0.122772 | 0.033427 |
| *Sa.* (*Sab.*) *purpureus* | 0.756885 | 0.045597 | -0.010218 | -0.097900 | -0.632555 | 0.087167 | 0.087369 |
| *Ae.* (*Och.*) *scapularis* | -0.755781 | -0.595029 | 0.202372 | 0.101374 | -0.101497 | 0.105245 | 0.046117 |
| *Ae.* (*Stg.*) *aegypti* | -0.740113 | -0.618213 | 0.185865 | 0.096589 | -0.103972 | 0.113762 | 0.049186 |
| *Sa.* (*Sbo.*) *chloropterus* | 0.738539 | -0.328664 | 0.200608 | -0.078782 | 0.502638 | 0.207241 | -0.067052 |
| *Ps.* (*Jan.*) *albigenu* | 0.669787 | -0.194045 | 0.359696 | 0.581486 | -0.110656 | 0.181694 | -0.031096 |
| *Sa.* (*Sab.*) *belisarioi* | 0.574237 | -0.218777 | 0.517919 | 0.377660 | -0.221176 | -0.399166 | 0.057180 |
| *Wy.* (*Tra.*) *aporonoma* | -0.204098 | 0.835920 | 0.160076 | 0.338607 | 0.322828 | 0.120343 | 0.024552 |
| *Ps.* (*Jan.*) *amazonica* | 0.342815 | 0.803056 | -0.329723 | -0.041891 | -0.280803 | 0.205066 | -0.078767 |
| *Hg.* (*Con.*) *leucocelaenus* | -0.214107 | 0.798123 | -0.539033 | 0.046284 | -0.082088 | -0.080013 | 0.106393 |
| *Tr. digitatum* | -0.234856 | 0.693925 | 0.316784 | 0.530398 | -0.175108 | -0.223105 | -0.034600 |
| *Sa.* (*Sab.*) *albiprivus* | -0.090981 | 0.538898 | 0.523317 | -0.446658 | -0.275549 | 0.388074 | -0.037664 |
| *Wy.* (*Den.*) *ypsipola* | -0.283637 | 0.424983 | 0.793152 | -0.240676 | 0.154689 | -0.125883 | -0.110224 |
| *Hg.* (*Haemagogus*) *janthinomys* | 0.259890 | -0.052911 | -0.702235 | -0.578506 | 0.059604 | -0.313242 | -0.013475 |
| *Ae.* (*Ochlerotatus*) *hastatus* | -0.181636 | 0.556598 | 0.692972 | -0.240816 | 0.001840 | -0.344694 | 0.013680 |
| *Ae.* (*Ochlerotatus*) *nubilus* | -0.265374 | 0.505794 | -0.397601 | 0.656519 | 0.170531 | 0.105561 | -0.210764 |
| *Cx.* (*Culex*) *coronator* | -0.415066 | 0.356325 | -0.481745 | 0.615289 | 0.296293 | 0.041815 | 0.023582 |
| *Ps.* (*Janthinosoma*) *circumflava* | 0.580104 | -0.285819 | 0.179727 | 0.606811 | -0.230730 | -0.336269 | 0.122279 |
| *Sa.* (*Sabethes*) *tarsopus* | 0.011746 | 0.526111 | 0.540802 | -0.547797 | 0.245682 | -0.204447 | 0.168411 |
| *Wy.* (*Cruzmyia*) *kummi* | 0.096084 | 0.216506 | -0.251470 | -0.294406 | -0.749935 | 0.418944 | 0.236781 |
| *Sa.* (*Sabethoides*) *tridentatus* | 0.348467 | -0.099786 | 0.199921 | -0.242388 | 0.647862 | 0.587949 | -0.066961 |
| *Cx.* (*Melanoconion*) *innovator* | 0.056862 | -0.059038 | -0.378357 | -0.223962 | 0.461347 | -0.284329 | 0.711536 |
| *Sa.* (*Sabethes*) *lanei* | 0.092382 | -0.182097 | -0.363347 | -0.421768 | -0.003900 | -0.392021 | -0.703350 |

Shaded cells have absolute loading values < ±0.3.

**Supplementary Table 3.** Species diversity

**Supplementary Table 3.** Species richness, diversity, and evenness by edge type, and by edge type and season at respective heights. Arithmetic means (± 1 standard error) calculated per sampling site (N = 3).

*By edge type*

| Edge type | N = | Richness | Diversity (*H'*) | Evenness |
|---|---|---|---|---|
| Continuous | 3 | 40.0 (0.58) | 2.60 (0.07) | 0.71 (0.02) |
| Treefall | 3 | 36.0 (3.06) | 2.10 (0.15) | 0.59 (0.03) |
| Rural | 3 | 33.3 (0.33) | 2.90 (0.06) | 0.83 (0.02) |
| Urban | 3 | 21.7 (5.78) | 1.98 (0.08) | 0.67 (0.05) |

*H'* = Shannon-Wiener diversity index.

*By edge type and season at 0 m*

| Edge type | N = | Richness | | Diversity (*H'*) | | Evenness | |
|---|---|---|---|---|---|---|---|
| | | Rainy | Dry | Rainy | Dry | Rainy | Dry |
| Continuous | 3 | 25.3 (1.45) | 24.0 (2.52) | 2.57 (0.07) | 2.67 (0.10) | 0.80 (0.01) | 0.84 (0.01) |
| Treefall | 3 | 25.0 (1.00) | 20.3 (3.76) | 2.13 (0.17) | 2.23 (0.26) | 0.66 (0.04) | 0.74 (0.04) |
| Rural | 3 | 24.0 (2.31) | 19.0 (2.52) | 2.59 (0.07) | 2.59 (0.21) | 0.82 (0.02) | 0.88 (0.04) |
| Urban | 3 | 15.7 (5.17) | 11.0 (3.51) | 1.64 (0.02) | 1.64 (0.18) | 0.63 (0.06) | 0.72 (0.02) |

*H'* = Shannon-Wiener diversity index.

*By edge type* and season at 5m*

| Edge type | N = | Richness | | Diversity (*H'*) | | Evenness | |
|---|---|---|---|---|---|---|---|
| | | Rainy | Dry | Rainy | Dry | Rainy | Dry |
| Continuous | 3 | 18.7 (0.88) | 14.0 (1.15) | 1.83 (0.17) | 1.90 (0.14) | 0.62 (0.06) | 0.72 (0.03) |
| Treefall | 3 | 16.7 (1.86) | 17.0 (2.31) | 1.46 (0.08) | 1.87 (0.22) | 0.52 (0.02) | 0.66 (0.06) |
| Rural | 3 | 11.0 (3.61) | 9.70 (2.85) | 1.93 (0.31) | 1.80 (0.28) | 0.88 (0.04) | 0.85 (0.03) |

*H'* = Shannon-Wiener diversity index.

*Urban edge excluded due to small sample size.

*The following pages show results of statistical testing for relevant comparisons of* ***species evenness****.*

**Supplementary Table 3.** Species diversity

*By edge type*

Results of one-way ANOVA to compare evenness by edge type, and Tukey HSD post-hoc tests. DF = degrees of freedom; significant P values highlighted in bold.

| **Comparison** | **DF =** | **One-way ANOVA** | **Pairwise comparison** | **Tukey HSD, P =** |
|---|---|---|---|---|
| Edge type | 3 | F = 9.46, P = 0.005 | Rural*, Treefall | **0.0036** |
| | | | Rural*, Urban | **0.0328** |
| | | | Continuous, Treefall | 0.1136 |
| | | | Rural, Continuous | 0.1183 |
| | | | Urban, Treefall | 0.3797 |
| | | | Continuous, Urban | 0.7988 |

*Significantly higher species evenness.

**Supplementary Table 3.** Species diversity

*By edge type and height*

Results of two-way ANOVA to compare evenness by edge type and height. DF = degrees of freedom; significant P values highlighted in bold.

| **Comparison** | **DF =** | **Two-way ANOVA** |
|---|---|---|
| Edge type | 3 | F = 10.6, **P = 0.0004** |
| Height | 1 | F = 0.11, P = 0.7452 |
| Edge type*height | 3 | F = 21.8, **P < 0.0001** |

Results of one-way ANOVA to compare evenness by edge type at 0 m, and Tukey HSD post-hoc tests. DF = degrees of freedom; significant P values highlighted in bold.

| **Comparison** | **DF =** | **One-way ANOVA** | **Pairwise comparison** | **Tukey HSD, P =** |
|---|---|---|---|---|
| Edge type 0 m | 3 | F = 9.78, **P = 0.0047** | Rural*, Treefall | **0.0292** |
| | | | Rural*, Urban | **0.0072** |
| | | | Continuous, Treefall | 0.0899 |
| | | | Rural, Continuous | 0.8517 |
| | | | Urban, Treefall | 0.7262 |
| | | | Continuous*, Urban | **0.0207** |

*Significantly higher species evenness.

Results of one-way ANOVA to compare evenness by edge type at 5 m, and Tukey HSD post-hoc tests. DF = degrees of freedom; significant P values highlighted in bold.

| **Comparison** | **DF =** | **One-way ANOVA** | **Pairwise comparison** | **Tukey HSD, P =** |
|---|---|---|---|---|
| Edge type 5 m | 3 | F = 33.5, **P < 0.0001** | Rural*, Treefall | **0.0005** |
| | | | Rural, Urban | 0.3270 |
| | | | Continuous, Treefall | 0.1522 |
| | | | Rural*, Continuous | **0.0073** |
| | | | Urban*, Treefall | **< 0.0001** |
| | | | Continuous, Urban* | **0.0009** |

*Significantly higher species evenness.

**Supplementary Table 3.** Species diversity

*By edge type and season at 0 m*

Results of two-way ANOVA to compare evenness by edge type and season. DF = degrees of freedom; significant P values highlighted in bold.

| Comparison | DF = | Two-way ANOVA |
|---|---|---|
| Edge type | 3 | F = 11.8, **P = 0.0003** |
| Season | 1 | F = 7.49, **P = 0.0147** |
| Edge type*season | 3 | F = 0.12, P = 0.9442 |

Results of a least-squares (LS) means Tukey HSD post-hoc test to compare evenness at forest edges. Edges not connected by the same letter are significantly different.

| Edge type | LS mean | Connecting letter | |
|---|---|---|---|
| Rural | 0.849 | A | |
| Continuous | 0.819 | A | |
| Treefall | 0.702 | | B |
| Urban | 0.674 | | B |

Results of a least-squares (LS) means Student's t-test post-hoc test to compare evenness between seasons. Seasons not connected by the same letter are significantly different.

| Season | LS mean | Connecting letter | |
|---|---|---|---|
| Dry | 0.795 | A | |
| Rainy | 0.727 | | B |

**Supplementary Table 3.** Species diversity

*By edge type and season at 5m*

Results of two-way ANOVA to compare evenness by edge type and season. DF = degrees of freedom; significant P values highlighted in bold.

| **Comparison** | **DF =** | **Two-way ANOVA** |
|---|---|---|
| Edge type | 2 | F = 21.6, **P = 0.0001** |
| Season | 1 | F = 3.98, P = 0.0691 |
| Edge type*season | 2 | F = 2.12, P = 0.1624 |

Results of a least-squares (LS) means Tukey HSD post-hoc test to compare evenness at forest edges. Edges not connected by the same letter are significantly different.

| **Edge type** | **LS mean** | **Connecting letter** | |
|---|---|---|---|
| Rural | 0.863 | A | |
| Continuous | 0.673 | | B |
| Treefall | 0.590 | | B |

Results of a least-squares (LS) means Student's t-test post-hoc test to compare evenness between seasons. Seasons not connected by the same letter are significantly different.

| **Season** | **LS mean** | **Connecting letter** |
|---|---|---|
| Dry | 0.743 | A |
| Rainy | 0.674 | A |

**Supplementary Table 4.** Rainy season vs. dry season abundance

**Supplementary Table 4.** Differences in mosquito abundance by forest edge type and season based on the eight most abundant species. Comparisons with fewer than 60 mosquitoes were excluded.

| | | Rainy season | | | | | Dry season | | | | | Wilcoxon Rank Sum | | |
|---|---|---|---|---|---|---|---|---|---|---|---|---|---|---|
| **Species** | **Edge type** | **$N^{mos}$ (d)** | **Min** | **Med** | **Max** | **IQR** | **$N^{mos}$ (d)** | **Min** | **Med** | **Max** | **IQR** | **DF** | **$\chi^2$** | **P =** |
| *Hg. janthinomys* | Continuous | 273 (18) | 0 | 13 | 61 | 15.8 | 159 (18) | 1 | 9 | 28 | 11.5 | 1 | 2.13 | 0.14 |
| *Hg. janthinomys* | Treefall | 490 (18) | 2 | 26.5 | 54 | 26.5 | 222 (19) | 1 | 8 | 75 | 10 | 1 | 12.5 | 0.0004 |
| *Ps. amazonica* | Continuous | 210 (18) | 0 | 2.5 | 42 | 21.5 | 98 (18) | 0 | 0 | 35 | 5 | 1 | 2.33 | 0.13 |
| *Ps. amazonica* | Treefall | 298 (18) | 0 | 4.5 | 85 | 17.3 | 101 (19) | 0 | 0 | 37 | 3 | 1 | 3.44 | 0.06 |
| *Ps. amazonica* | Rural | 94 (17) | 0 | 0 | 28 | 12.5 | 25 (18) | 0 | 0 | 20 | 0.25 | 1 | 2.95 | 0.086 |
| *Li. durhamii* | Rural | 50 (17) | 0 | 4 | 6 | 3 | 16 (18) | 0 | 1 | 3 | 1 | 1 | 9.43 | 0.002 |
| *Li. durhamii* | Urban | 192 (18) | 0 | 1.5 | 64 | 11.5 | 58 (20) | 0 | 0.5 | 23 | 4.75 | 1 | 2.21 | 0.14 |
| *Sa. chloropterus* | Continuous | 45 (18) | 0 | 2 | 9 | 3.25 | 42 (18) | 0 | 1.5 | 7 | 3.5 | 1 | 0.04 | 0.83 |
| *Sa. chloropterus* | Treefall | 52 (18) | 0 | 2.5 | 7 | 3.25 | 38 (19) | 0 | 1 | 13 | 3 | 1 | 3.84 | 0.0499 |
| *Sa. chloropterus* | Rural | 32 (17) | 0 | 1 | 6 | 2 | 34 (18) | 0 | 1 | 7 | 3.25 | 1 | 0.7 | 0.4 |
| *Wy. aporonoma* | Continuous | 46 (18) | 0 | 2.5 | 6 | 3.25 | 27 (18) | 0 | 1 | 4 | 3 | 1 | 2.51 | 0.11 |
| *Ae. albopictus* | Urban | 71 (18) | 0 | 3 | 12 | 3.75 | 46 (20) | 0 | 1.5 | 7 | 2 | 1 | 2.96 | 0.085 |
| *Li. pseudomethysticus* | Continuous | 46 (18) | 0 | 2 | 11 | 3.25 | 22 (18) | 0 | 0 | 8 | 1.5 | 1 | 3.72 | 0.054 |

**$N^{mos}$ (d)** = number of mosquitoes sampled (number of sampling days), **Min** = minimum, **Med** = median, **Max** = maximum, **IQR** = interquartile range, **DF** = degrees of freedom, **$\chi^2$** = chi-square value, **P =** = P value. Gray cells show significant P values.

**Supplementary Table 5.** Species occurrence

**Supplementary Table 5.** Occurrence of the eight most abundant species overall showing the number (N =) and percent (%) days when a species was encountered across both heights. Total 71 rainy season sampling days across edge types at 0 m and 5 m = 142 Y/N outcomes; 75 dry season sampling days across edge types at 0 m and 5 m = 150 Y/N outcomes.

| **Species** | **Total** N = (%) | **Continuous** N = (%) | **Treefall** N = (%) | **Rural** N = (%) | **Urban** N = (%) |
|---|---|---|---|---|---|
| *RAINY SEASON* | | | | | |
| *Hg. janthinomys* | 78 (54.9) | 29 (80.6) | 35 (97.2) | 11 (32.4) | 3 (8.3) |
| *Ps. amazonica* | 64 (45.1) | 20 (55.6) | 22 (61.1) | 15 (44.1) | 7 (19.4) |
| *Li. durhamii* | 37 (26.1) | 6 (16.7) | 2 (5.6) | 15 (44.1) | 14 (38.9) |
| *Sa. chloropterus* | 66 (46.5) | 16 (44.4) | 25 (69.4) | 18 (52.9) | 7 (19.4) |
| *Wy. aporonoma* | 48 (33.8) | 20 (55.6) | 11 (30.6) | 13 (38.2) | 4 (11.1) |
| *Ae. albopictus* | 29 (20.4) | 1 (2.8) | 0 (0) | 8 (23.5) | 20 (55.6) |
| *Sa. cyaneus* | 36 (25.4) | 14 (38.9) | 14 (38.9) | 7 (20.6) | 1 (2.8) |
| *Li. pseudomethysticus* | 29 (20.4) | 13 (36.1) | 10 (27.8) | 5 (14.7) | 1 (2.8) |
| *DRY SEASON* | | | | | |
| *Hg. janthinomys* | 74 (49.3) | 27 (75.0) | 35 (92.1) | 10 (27.8) | 2 (5.0) |
| *Ps. amazonica* | 34 (22.7) | 13 (36.1) | 13 (34.2) | 5 (13.9) | 3 (7.5) |
| *Li. durhamii* | 28 (18.7) | 2 (5.6) | 2 (5.3) | 13 (36.1) | 11 (27.5) |
| *Sa. chloropterus* | 45 (30.0) | 15 (41.7) | 14 (36.8) | 13 (36.1) | 3 (7.5) |
| *Wy. aporonoma* | 35 (23.3) | 14 (38.9) | 9 (23.7) | 10 (27.8) | 2 (5.0) |
| *Ae. albopictus* | 28 (18.7) | 2 (5.6) | 0 (0) | 8 (22.2) | 18 (45.0) |
| *Sa. cyaneus* | 38 (25.3) | 17 (47.2) | 11 (28.9) | 8 (22.2) | 2 (5.0) |
| *Li. pseudomethysticus* | 14 (9.3) | 7 (19.4) | 3 (7.9) | 4 (11.1) | 0 (0) |

**Supplementary Table 6.** Standard least squares analysis

**Supplementary Table 6.** Standard least squares analysis (P value and (F Ratio)) testing associations between species abundance and environmental variables in the rainy season. Blue and red shaded cells indicate significant positive or negative associations, respectively.

| | | | | | | *7-day cumulative rainfall lag* | | | |
|---|---|---|---|---|---|---|---|---|---|
| **Species [N=]*** | **Edge type**** | **Height†** | **Mean temp** | **Mean RH** | **Mean weather‡** | **1 wk** | **2 wk** | **3 wk** | **4 wk** |
| | | | | *RAINY SEASON* | | | | | |
| *Hg. janthinomys* [1,144] | C, T | <0.0001 (27.0) | <0.0001 (34.8) | | | | | | |
| *Ps. amazonica* [826] | C, T, R | | | | | | | 0.0005 (12.8) | <0.0001 (24.9) |
| *Li. durhamii* [316] | R, U | 0.002 (10.8) | | | | | | | |
| *Sa. chloropterus* [243] | C, T, R | <0.0001 (42.1) | | | 0.002 (10.1) | | | | |
| *Wy. aporonoma* [131] | C, T, R | 0.002 (10.5) | | | | | | | |
| *Ae. albopictus* [117] | U | 0.0002 (17.4) | | | | | | | |
| *Sa. cyaneus* [110] | C, T, R | | | | 0.002 (9.6) | 0.03 (4.6) | | | |
| *Li. pseudomethy-sticus* [110] | C, T, R | <0.0001 (25.0) | | | | | | | |

*Number of mosquitoes sampled across edge types included in the analysis.
**Edge type included in the analysis: C = continuous forest, T = treefall gap, R = rural, U = urban. Note that edge type was not included as a variable.
†Red and blue shaded cells indicate that a species was more common at ground level or on 5 m platforms, respectively.
‡Red shaded cells indicate a negative association with mean weather i.e., an increase in abundance during clearer conditions.

**Supplementary Table 7.** Generalized linear model

**Supplementary Table 7.** Generalized linear model testing effects of edge type and height on the rainy and dry season occurrence of the eight most abundant species overall. Cells shaded in light gray indicate marginally significant P values (0.01 > P < 0.1), while those in dark gray show unreported P values due to highly significant (P < 0.01) interaction effects.

| Species | Height | Edge type | Edge type x height | Model P | Model $\chi^2$ |
|---|---|---|---|---|---|
| *RAINY SEASON* | | | | | |
| *Hg. janthinomys* | 0.04 | <0.0001 | 0.55 | <0.0001 | 45.1 |
| *Ps. amazonica* | 0.04 | 0.0004 | 0.61 | 0.0004 | 26.6 |
| *Li. durhamii** | - | - | <0.0001 | <0.0001 | 47.9 |
| *Sa. chloropterus* | 0.002 | 0.006 | 0.04 | 0.0003 | 26.9 |
| *Wy. aporonoma* | 0.01 | 0.007 | 0.85 | 0.019 | 16.7 |
| *Ae. albopictus** | - | - | <0.0001 | <0.0001 | 59.0 |
| *Sa. cyaneus* | 0.92 | 0.001 | 0.11 | 0.003 | 21.9 |
| *Li. pseudomethysticus** | - | - | <0.0001 | <0.0001 | 66.0 |
| *DRY SEASON* | | | | | |
| *Hg. janthinomys* | - | - | <0.0001 | <0.0001 | 66.7 |
| *Ps. amazonica* | 0.005 | 0.0005 | 0.55 | <0.0001 | 30.6 |
| *Li. durhamii** | - | - | 0.002 | <0.0001 | 31.7 |
| *Sa. chloropterus* | - | - | 0.003 | <0.0001 | 35.7 |
| *Wy. aporonoma* | 0.004 | 0.02 | 0.69 | 0.017 | 17.0 |
| *Ae. albopictus** | - | - | <0.0001 | <0.0001 | 58.3 |
| *Sa. cyaneus* | 0.76 | 0.008 | 0.33 | 0.008 | 11.2 |
| *Li. pseudomethysticus* | <0.0001 | 0.0004 | 0.012 | 0.0002 | 28.1 |

*Highly significant interaction effects not considered due to small number of mosquitoes sampled above ground throughout the study (N ≤ 6).

**Supplementary Table 8.** Contingency table analysis

**Supplementary Table 8.** Contingency table analysis and Pearson's chi-square test of *Sabethes* mosquitoes based on 30-minute occurrence (Y/N) data grouped at subgenus level. DF = 1 for all comparisons.

*By height*

| Subgenus | N =* | Height (m) | Y | N | $\chi^2$ | P = |
|---|---|---|---|---|---|---|
| *Sabethes* | 354 | 0 | 160 | 1300 | 3.72 | 0.054 |
| | | 5 | 194 | 1266 | | |
| *Sabethoides* | 257 | 0 | 41 | 1419 | 131 | <0.0001 |
| | | 5 | 216 | 1244 | | |

*Number of 30-minute intervals during which subgenus was sampled.

*By edge type and height*

| Edge type | Subgenus | N =* | Height (m) | Y | N | $\chi^2$ | P = |
|---|---|---|---|---|---|---|---|
| Continuous | *Sabethes* | 100 | 0 | 31 | 329 | 16.8 | <0.0001 |
| | | | 5 | 69 | 291 | | |
| | *Sabethoides* | 87 | 0 | 5 | 355 | 77.5 | <0.0001 |
| | | | 5 | 82 | 278 | | |
| Treefall | *Sabethes* | 136 | 0 | 77 | 293 | 2.92 | 0.1 |
| | | | 5 | 59 | 311 | | |
| | *Sabethoides* | 80 | 0 | 12 | 358 | 44 | <0.0001 |
| | | | 5 | 68 | 302 | | |
| Rural | *Sabethes* | 81 | 0 | 37 | 313 | 0.68 | 0.4 |
| | | | 5 | 44 | 306 | | |
| | *Sabethoides* | 74 | 0 | 17 | 333 | 24.2 | <0.0001 |
| | | | 5 | 57 | 293 | | |
| Urban | *Sabethes* | 37 | 0 | 15 | 365 | 1.39 | 0.2 |
| | | | 5 | 22 | 358 | | |
| | *Sabethoides* | 16 | 0 | 7 | 373 | 0.26 | 0.6 |
| | | | 5 | 9 | 371 | | |

*Number of 30-minute intervals during which subgenus was sampled.
